# Moving Targets: Monitoring Target Trends in Drug Discovery by Mapping Targets, GO Terms, and Diseases

**DOI:** 10.1101/691550

**Authors:** Barbara Zdrazil, Lars Richter, Nathan Brown, Rajarshi Guha

## Abstract

Drug Discovery is a lengthy and costly process and has faced a period of declining productivity within the last two decades. As a consequence, integrative data-driven approaches are nowadays on the rise in pharmaceutical research, making use of an inter-connected (network) view on diseases. In addition, evidence-based decisions are alleviated by studying the time evolution of innovation trends in drug discovery.

In this paper a new approach leveraging data mining and data integration for inspecting target innovation trends protein family-wise is presented. The study highlights protein families which are receiving emerging interest in the drug discovery community (mainly kinases and G protein coupled receptors) and those with areas of interest in target space that have just emerged in the scientific literature (mainly kinases and transporters) highlighting novel opportunities for drug intervention.

In order to delineate the evolution of target-driven research interest from a biological perspective, trends in biological process annotations from Gene Ontology (GO) and disease annotations from DisGeNet for major target families are captured. The analysis reveals an increasing interest in targets related to immune system processes, and a recurrent trend for targets involved in circulatory system processes. At the level of disease annotations, targets associated to e.g., cancer-related pathologies as well as to intellectual disability and schizophrenia are increasingly investigated nowadays.

Can this knowledge be used to study the “movement of targets” in a network view and unravel new links between diseases and biological processes? We tackled this question by creating dynamic network representations considering data from different time periods. The dynamic network for immune system process-associated targets suggest that e.g. breast cancer as well as schizophrenia are linked to the same targets (cannabinoid receptor CB2 and VEGFR2) thus suggesting similar treatment options which could be confirmed by literature search. The methodology has the potential to identify other drug repurposing candidates and enables researchers to capture trends in research attention in target space at an early stage.

The KNIME workflows and R scripts used in this study are publicly available from https://github.com/BZdrazil/Moving_Targets.

**Author summary:** In this study we have investigated target innovation in drug discovery over a period of 22 years (1995-2016) by extracting time trends of research interest (as published in the scientific literature and stored in the ChEMBL database) in certain protein classes inspecting different measures (numbers of pharmacological measurements, targets, papers, and drugs). Focusing on the most relevant protein classes in drug discovery (G protein-coupled receptors, kinases, ion channels, nuclear receptors, proteases, and transporters), we further linked single targets to Gene Ontology (GO) biological process annotations and inspected steep increasing or decreasing trends of GO annotations within target families over time. We also tracked trends in disease annotations from DisGeNET by filtering out diseases linked to targets with emerging trends in research interest. Finally, targets, GO terms, and diseases are interconnected in network representations and shifts in research foci are investigated over time. This new methodology which utilizes data mapping and data analysis can be used to explore trends in research attention target family-wise, to uncover previously unknown links between diseases and biological processes and to identify potential candidates for drug repurposing.

## Introduction

The organization of proteins (and genes) into families, is usually achieved through phylogeny (amino acid sequence similarity) and - to a lesser extent - by similarity of secondary structures. The assumption is that proteins belonging to the same family share a common function which will also be encoded in the protein sequences and their folds.

Apart from the general desire to organize information, such indexing and grouping of proteins is especially relevant in drug discovery. If we understand how protein modulation, e.g. by small molecules works for a particular protein, we also move closer towards understanding how modulating a whole group of related proteins might work. Thus, structure-function relationships are often carried out at the protein family level [1, 2].Somewhat contradictory to this observation is the phenomenon of polypharmacology, where a molecule interacts with multiple (often unrelated) targets (either wanted or unwanted) [3–5].Therefore, across target families, close similarities in binding patterns appear to occur for similar ligands - this has been explored systematically for the identification of novel ligand-target associations [6–8]. A straightforward application of exploring polypharmacology of drugs is for repurposing reasons [9], especially in the area of rare diseases [10].

Setting aside the goal of unraveling ligand-protein interactions, another emerging application of target (family) indexing and characterization in drug discovery is the desire to capture the past, current, and (potential) future degree of research interest in main drug target families. Until now, exploratory studies of research attention across target families have been carried out mainly from a static point of view with only a few analyses including the time dimension. In an analysis article from 2011 by Rask-Andersen and colleagues [11], the authors looked into such trends as the exploitation of new drug targets over a period of 30 years and have identified three innovation peaks. Interestingly, when exploring drug-target networks their analysis revealed that newer molecular entities tend to form isolated networks pointing towards a tendency to interact with new drug targets and potentially possessing new modalities for treatment.

In a more recent analysis by Santos et al. [12] the authors highlight the main drug efficacy target families, which together account for approximately 44% of all human drug targets: G-protein coupled receptors (GPCRs), kinases, ion channels, and nuclear receptors. The authors also present a temporal view on innovation patterns in these privileged protein classes by analyzing the proportion of approved drugs for each target class comparing different time frames. In addition, patterns of innovation in different therapeutic areas are represented in a timewise manner, but independent of the target class.

Since 2014, a strategic effort by the US National Institutes of Health termed the Illuminating the Druggable Genome (IDG) initiative focuses on exploring currently understudied, but potentially druggable, proteins [13]. In order to be able to define the degree to which a target is comparatively well studied or understudied, the existing knowledge on current associations between compounds (or drugs), targets, Mechanism of Action (MoA), and disease were systematically collected and mapped by this initiative. These efforts resulted in defining so-called Target Developmental Level (TDL) categories by which targets can be classified according to the depth of investigations undertaken for a particular target (by uniting available chemical, biological and clinical information). The four categories in the TDL scheme, Tclin (clinic: drug annotation(s) with approved MoA), T_chem_ (chemistry: binding small molecules with high potency), T_bio_ (biology: disease annotation(s) available), and T_dark_ (dark genome: not falling into one of the other categories), are thereafter representing different (decreasing) stages of target innovation. A multimodal web interface called PHAROS enables intuitive access to the IDG knowledgebase [14]. A similar European initiative, called Open Targets, uses human genetics and genomics data for systematic drug target identification and prioritization [15].

Considering the compound space, investigating drug-target associations does provide only a limited picture of innovation trends in drug discovery. The stage at which NMEs are approved is the very final stage of a long series of investigations and decisions taken. Thus, including all data available at the compound level over time (including bioactivity data extracted from databases and from the scientific literature) is a crucial step forward in the domain of data-driven drug discovery strategies.

The immense pool of small compound data spans two main data repertoires where preclinical data is frequently reported: scientific literature and patents. A thorough analysis of trends in publishing behavior of bioactivity compounds for novel targets (rather literature or patents first) was delivered by Ashenden et al. [16]. Preclinical data can guide decision making at earlier stages in the drug discovery pipeline since it reflects trends in research attention within the community. It might help researchers to base their research endeavors on evidence-based criteria and direct their interest towards understudied proteins or core scaffolds [17].

In this study, major target classes are monitored over time to capture the different aspects of target innovation in drug discovery. First, target families of emerging interest in drug discovery are highlighted as reflected by an increasing number of targets, bioactivity measurements, annotated drugs, and number of publications. Second, trends in GO (Gene Ontology) biological process annotations enable to filter out biological processes of (potential) emerging attention per target family. Third, studying the evolution of disease annotations for major target families highlights popular targets of intervention for particular diseases. Finally, uniting the different views in the form of network representations by mapping targets, GO terms, and diseases, allows us to uncover new links between diseases and biological processes.

## Results and discussion

### Target innovation trends

We investigated protein innovation trends by inspecting different measures reflecting research attention per target family: increase or decrease of the number of unique compounds, number of drug-efficacy target annotations (playing a role in the efficacy of the drug in the indication(s) for which it is approved), number of published papers (as expressed by PMIDs), and number of unique targets. In addition, we studied the evolution of the proportions of Target Developmental Levels (TDLs) through the years for each target family.

For all four measures describing protein family popularity in absolute numbers (Figure 1), the steepest upward trends can be observed for GPCRs and kinases over 21 years (1995-2015). Interestingly, although the GPCR family was the most studied in terms of unique compounds and papers from the year 2000 onwards, kinases outpaced GPCRs towards the end of the observation period (in 2013 for compound counts, and in 2015 for paper counts). Investigating the trend lines delivered by target and drug-target annotation counts (Figure 1, bottom left and right), a less steady increase for the family of kinases can be observed since large fluctuations in these measures were taking place from 2005 onwards. The two largest peaks in terms of the number of published targets can be denoted for 2011 (359 unique targets) and 2008 (289 unique targets), which is not reflected in a particularly large number of papers for these years (106 and 90 papers, respectively). Looking a bit closer at the respective numbers of targets published in the respective papers of these years, it becomes clear that these two target peaks can each be attributed to one large study, which were both dedicated to comprehensive kinase inhibitor selectivity screens [18, 19]. As obvious from the steep rise in paper and compound counts from 2012/2013 onwards the two large scale screening studies (both designed by Zarrinkar et al.) might have sparked the increasing research interest in particularly this family of proteins since “the data set suggests compounds to use as tools to study kinases for which no dedicated inhibitors exist” [18]. In terms of numbers of drug-efficacy target annotations, a picture similar to that of target trends can be observed with also the highest peak in 2011 originating from the paper by Davis et al. (80 out of 90 total drug-target annotations in 2011).

**Figure 1.**
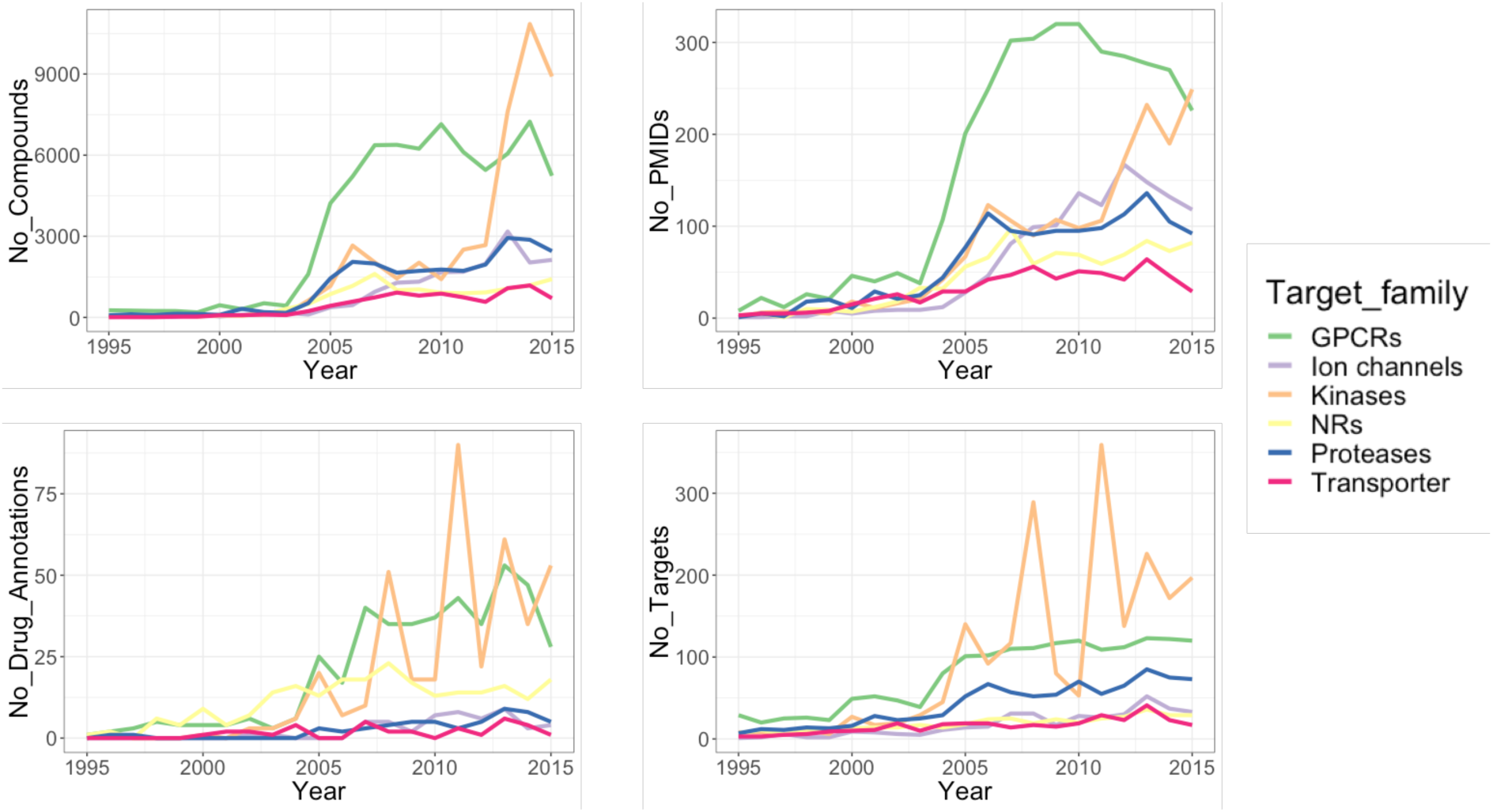
Graph showing the evolution of different measures of popularity for a respective target family over time in absolute numbers: numbers of unique compounds, numbers of Pubmed IDs, numbers of drug-efficacy target annotations, and numbers of targets.

In comparison to kinases, all other target families are showing relatively smooth increases in the different popularity measures depicted. Interestingly, GPCRs, although of emerging interest until recently, were obviously not so much affected by large screening studies (which is revealed by their high paper counts over the years and relatively flat compound and target trends between 2005 and 2015). Membrane proteins like GPCRs are extremely challenging targets when it comes to protein purification and crystallization. Therefore - although 33% of all small molecule drugs are targeting GPCRs - they account for only 12% of human protein drug targets [12].

From Figure 1 it also becomes visible that ion channels have outperformed proteases in terms of paper numbers over the years. This is not unexpected since ion channels according to Santos et al. make up the largest proportion of drug targets inspecting the individual protein families (19%) [12]. However, to date this is not so well reflected in terms of literature-reported compounds, targets and drug-efficacy target annotations (Figure 1).

By relating the absolute numbers from target innovation trends (as represented in Figure 1) to the respective total numbers of compounds, papers, drug-efficacy target annotations, and targets per year, relative trend measures are observed. These enable temporal comparison across the target families more easily, particularly when evaluating the early years of rational drug discovery, which tend to be relatively data poor. Notably, nuclear receptors outperformed all other target families in terms of drug-efficacy target annotations from 1998-2004, but not in terms of other popularity measures (Figure 2). This is also reflected by inspecting the proportions of small-molecule drugs and human drug targets, discussed in Santos et al. [12]: nuclear receptor drugs make up 16% of all drugs, but only 3% of drug targets. Inspecting the proportions of drug-efficacy target annotations over the whole observation period (Figure 2), it becomes clear that from 2005 onwards primarily GPCRs and later kinases display a relative enrichment in this popularity measure (in relation to drug-efficacy target annotations for all six target families). Notably, interest in kinases seems to be a more recent trend exemplified by the relative enrichment in all popularity measures under investigation by 2016 (Figure 2).

**Figure 2.**
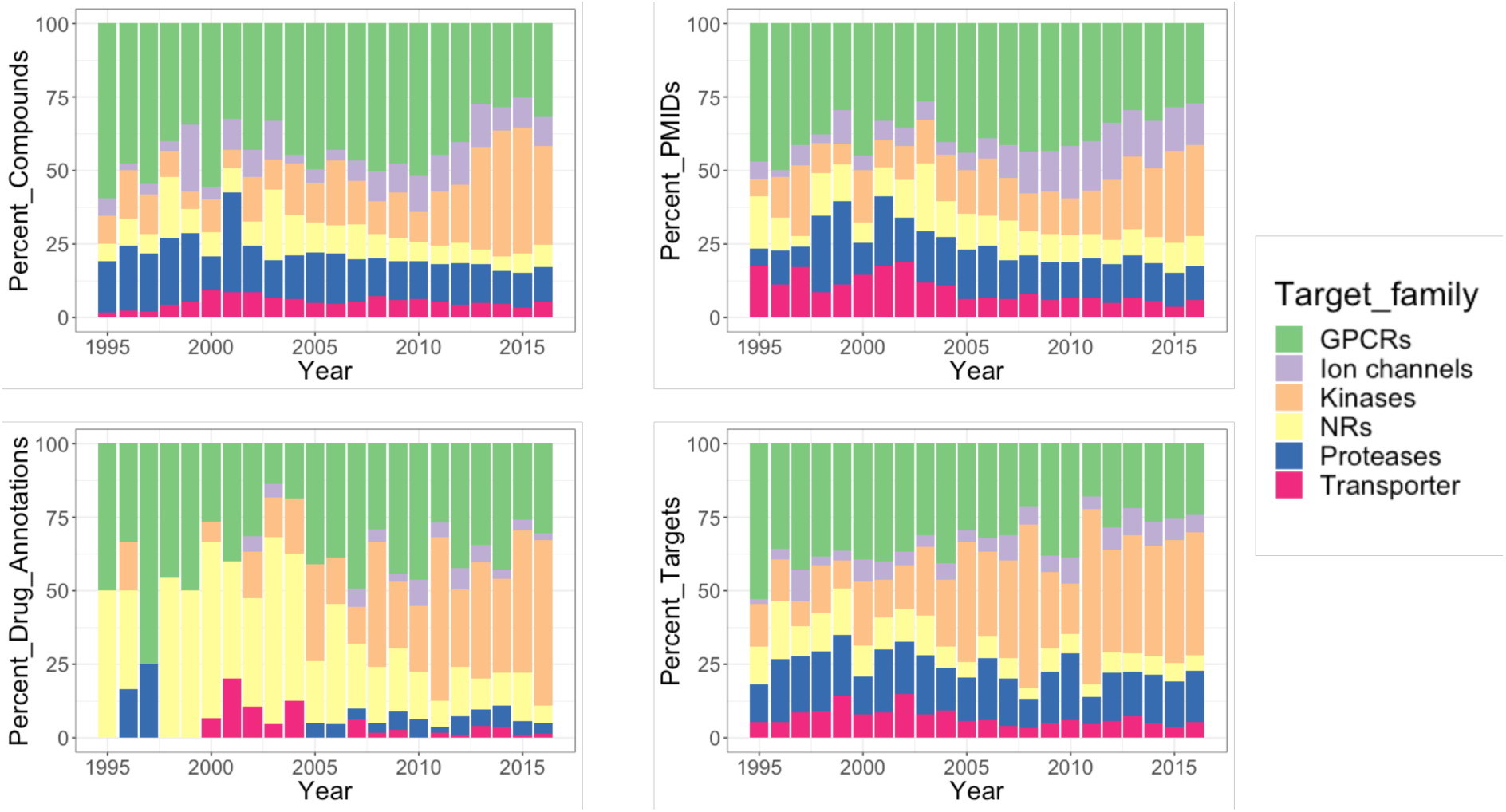
Bar charts showing the evolution of different measures of popularity for a respective target family over time expressed as proportions: percentages of compounds, percentages of Pubmed IDs, percentages of drug-efficacy target annotations, and percentages of targets.

A multi-dimensional measure of (evidence-based) research attention in particular human protein targets is provided by the Target Developmental Level (TDL) categories that were assigned by the IDG initiative [13, 14]. By looking at the time evolution of TDL categories protein family-wise (Figure 3), differences with respect to stages of drug development between the families can be captured. It has to be noted, that this visualization does not show the entirety of protein targets for each target family but reflects research endeavors, successes and future opportunities as reflected in the scientific literature (and recorded by ChEMBL). From Figure 3 it becomes obvious that within the Tclin category, ion channels and GPCRs make up the largest fraction of targets, consistently over the years. The majority of well-studied ion channel and GPCR proteins are already linked to at least one approved drug by MoA and proportionally not many new targets without known drugs (categories T_chem_ and T_bio_) have been reported in the scientific literature within the last ∼10 years. Inspecting the nuclear receptor superfamily the same tendency holds true for 1995-2005 but later the proportion of targets belonging to T_chem_ continuously increased, pointing to investigations on novel targets with potent compounds (but lacking approved drugs), such as investigations on metabolic nuclear receptors [20, 21].

**Figure 3.**
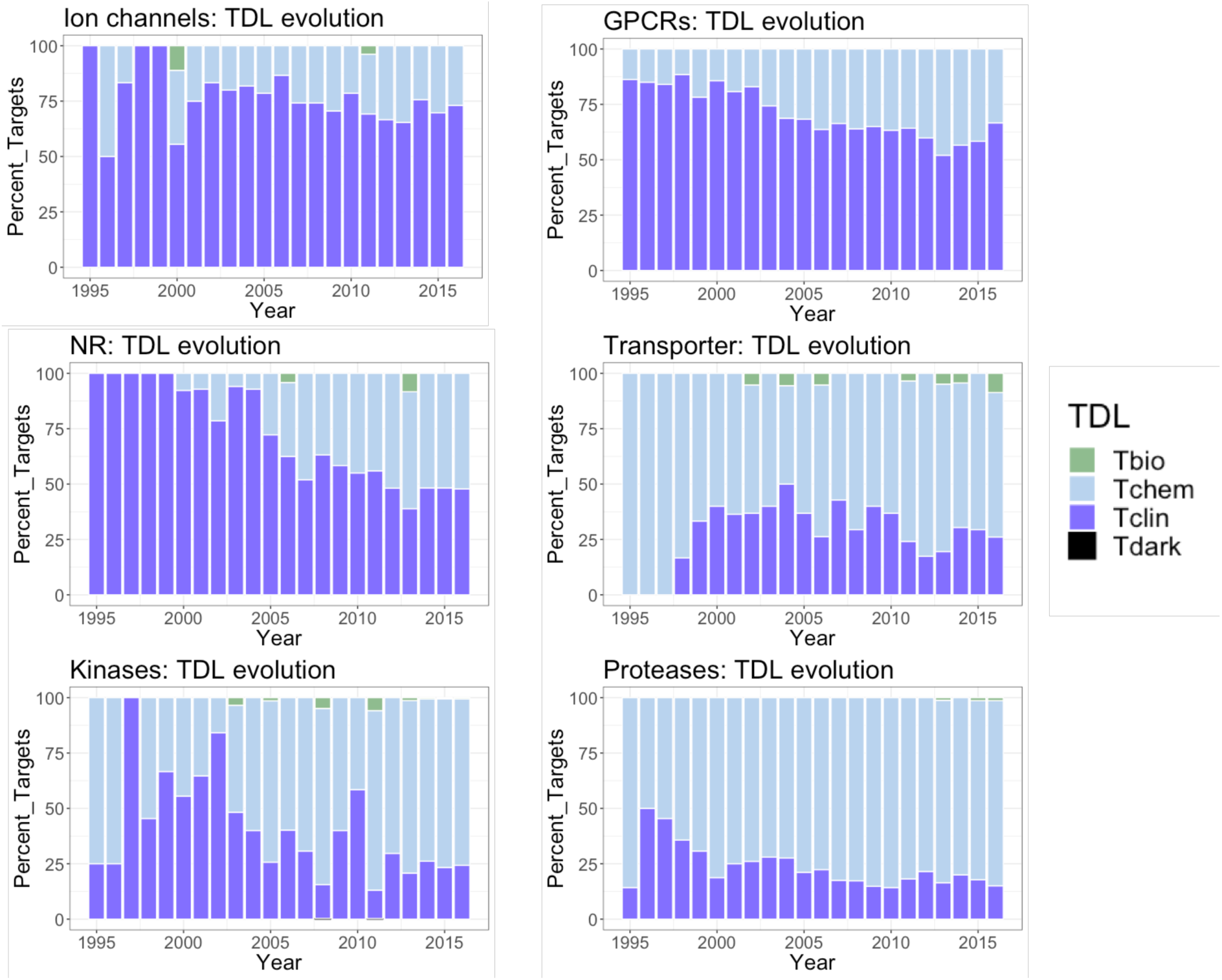
Bar charts showing the time evolution of the Target Developmental Level (TDL) categories across different target families.

In general, a higher fraction of targets from the categories T_chem_ or T_bio_ in recent years, highlights opportunities for drug discovery that are underexplored. These tendencies are more pronounced for the protein classes of transporters, kinases, and proteases (see Figure 3). For kinases, even targets of the category T_dark_ in 2008 and 2011 (one target each) are observed. The highest fractions of T_chem_ and T_bio_ (compared to all other years) can also be attributed to the same years which is the result of two large screening studies described above [18, 19].

Taken together the results from studying the evolution of target trend measures, a shift from GPCR-focused to kinase-focused research can be observed within the last two decades. Results from tracking the target developmental levels suggest that future research be directed towards unexploited areas in kinase, protease, transporter, and nuclear receptor focused studies since these families are still offering a wealth of targets with no approved drugs. It will however depend on the available research infrastructure, collaborations, and funded projects, if the “hard targets” (e.g. transmembrane transporters) can also be systematically deorphanized in the future. Initiatives like the EU-funded project Resolute [22] which concentrates on deorphanizing Solute Carriers will hopefully pave the way for other projects uncovering the unexploited opportunities in the human proteome.

### Evolution of Gene Ontology annotations

Moving from the level of target families to the level of individual targets enables us to explore biological aspects of target innovation trends encoded in Gene Ontology (GO) annotations. Here, we focus on GO biological process terms as they “represent a specific objective that the organism is genetically programmed to achieve”. Thus, by investigating steep increasing or decreasing trends of certain GO biological process annotations within target families over time, we are able to delineate the evolution of target-driven research interest from a biological perspective. Such trends are identified by fitting a robust regression line to the time-evolution of bioactivity counts (counts of measurements) with a certain GO annotation. Bioactivity numbers are expressed as percentages of total bioactivity counts in order to equilibrate the influence of a general upward trend in the number of measurements over time and to filter out trends that are showing an effect at the level of all target families (not only within a certain target family).

Interestingly, when considering only significant trends (p <= 0.05) the families of kinases and ion channels are showing exclusively positive GO trends (kinases: 31; ion channels: 7), whereas for proteases and NRs exclusively negative GO trends were extracted (proteases: 15; nuclear receptors: 10;). While for transporters no significant trends could be extracted by fitting a robust regression, the family of GPCRs delivered both positive and negative GO biological process annotation trends, but with the majority being negative (2 positive, 14 negative trends; see Table 1 and Supplementary Figure S1). Strikingly, for GPCRs, the biological process GO annotation “immune system process” (definition: any process involved in the development or functioning of the immune system; see also Supplementary Table S1) shows by far the steepest increasing trend (measured as total bioactivity counts) with a peak in 2009 (approx. 14% of all bioactivity measurements). The identified trend underpins the increasing interest in GPCRs as possible modulators of the immune system, such as the regulation of T cell activation, homeostasis and function [23]. Through inspection of the individual contribution of certain target classes to this pronounced trend, the research interest in target classes over time being involved in a certain biological process (i.e., “immune system process”) is delivered. For GPCRs, the positive trend observed for the “immune system process” GO term is largely driven by an increase in investigations on proteins belonging to the classes of opioid receptors, cannabinoid receptors, and CC-chemokine receptors (Figure 4).

**Figure 4.**
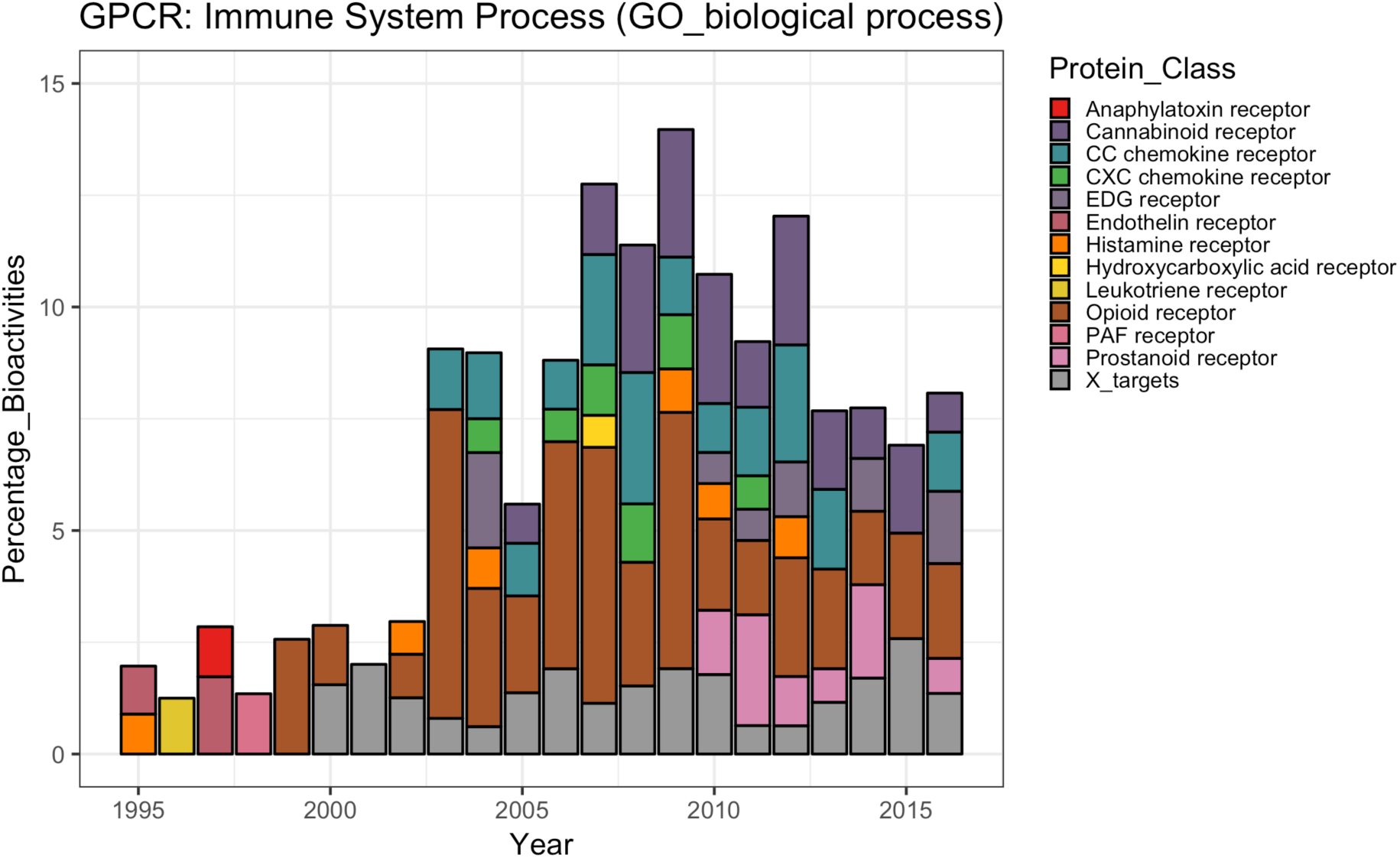
Bar chart showing the proportions of different protein classes contributing to the GO biological process annotation trend “immune system process” for GPCRs. Grey bars indicate target classes with little contribution to the general trend (less than 10% of the largest portion of a protein class in a certain year).

**Table 1.** List of annotated GO biological process annotations with statistically significant positive or negative trends (robust regression; p <= 0.05) for each target family. No significant trends could be observed for transporters.

| Target family | GO biological process annotation | Slope | Target family | GO biological process annotation | Slope |
| --- | --- | --- | --- | --- | --- |
| Kinase | phosphorylation | 1.606 | GPCR | immune system process | 0.440 |
| Kinase | cellular protein modification process | 1.605 | GPCR | locomotion | 0.247 |
| Kinase | signal transduction | 1.430 | GPCR | lipid metabolic process | -0.147 |
| Kinase | cell differentiation | 1.409 | GPCR | response to toxic substance | -0.158 |
| Kinase | multicellular organismal development | 1.122 | GPCR | digestion | -0.176 |
| Kinase | immune system process | 0.990 | GPCR | cell proliferation | -0.249 |
| Kinase | positive regulation of catalytic activity | 0.840 | GPCR | vesicle-mediated transport | -0.324 |
| Kinase | response to stress | 0.754 | GPCR | transport | -0.435 |
| Kinase | cell death | 0.663 | GPCR | metabolic process | -0.475 |
| Kinase | cell motility | 0.652 | GPCR | homeostatic process | -0.494 |
| Kinase | anatomical structure development | 0.573 | GPCR | response to stress | -0.536 |
| Kinase | cytoskeleton organization | 0.442 | GPCR | cell-cell signaling | -0.583 |
| Kinase | cell cycle | 0.400 | GPCR | signal transduction | -0.617 |
| Kinase | cell proliferation | 0.399 | GPCR | multicellular organismal development | -0.632 |
| Kinase | cell morphogenesis | 0.365 | GPCR | positive regulation of catalytic activity | -0.801 |
| Kinase | protein complex assembly | 0.363 | GPCR | circulatory system process | -1.235 |
| Kinase | reproduction | 0.318 | Protease | homeostatic process | -0.087 |
| Kinase | positive regulation of apoptotic process | 0.309 | Protease | female pregnancy | -0.109 |
| Kinase | anatomical structure formation involved in morphogenesis | 0.304 | Protease | cellular protein modification process | -0.117 |
| Kinase | homeostatic process | 0.287 | Protease | vesicle-mediated transport | -0.128 |
| Kinase | embryo development | 0.263 | Protease | symbiosis, encompassing mutualism through parasitism | -0.135 |
| Kinase | locomotion | 0.259 | Protease | embryo development | -0.138 |
| Kinase | biosynthetic process | 0.252 | Protease | multicellular organismal development | -0.139 |
| Kinase | symbiosis, encompassing mutualism through parasitism | 0.222 | Protease | anatomical structure formation involved in morphogenesis | -0.141 |
| Kinase | cellular nitrogen compound metabolic process | 0.220 | Protease | cell differentiation | -0.170 |
| Kinase | nucleocytoplasmic transport | 0.210 | Protease | immune system process | -0.251 |
| Kinase | protein targeting | 0.208 | Protease | response to stress | -0.264 |
| Kinase | transport | 0.204 | Protease | catabolic process | -0.281 |
| Kinase | vesicle-mediated transport | 0.201 | Protease | proteolysis | -0.292 |
| Kinase | aging | 0.161 | Protease | cell motility | -0.324 |
| Kinase | cell division | 0.138 | Protease | extracellular matrix organization | -0.514 |
| Ion channel | signal transduction | 0.259 | Nuclear receptor | female pregnancy | -0.100 |
| Ion channel | homeostatic process | 0.241 | Nuclear receptor | multicellular organismal development | -0.213 |
| Ion channel | transport | 0.211 | Nuclear receptor | reproduction | -0.220 |
| Ion channel | transmembrane transport | 0.211 | Nuclear receptor | positive regulation of apoptotic process | -0.226 |
| Ion channel | cellular protein modification process | 0.182 | Nuclear receptor | anatomical structure development | -0.244 |
| Ion channel | phosphorylation | 0.179 | Nuclear receptor | cell differentiation | -0.249 |
| Ion channel | response to stress | 0.126 | Nuclear receptor | biosynthetic process | -0.301 |
|  |  |  | Nuclear receptor | cellular nitrogen compound metabolic process | -0.301 |
|  |  |  | Nuclear receptor | signal transduction | -0.302 |
|  |  |  | Nuclear receptor | growth | -0.308 |

Investigations on opioid receptors have shown that their role in immune function can be manifold, ranging from immunosuppression, immunomodulation, and dual effects [24]. Opioid abuse itself can accelerate the progression of HIV infection, which is believed to happen due to changes of the immune status [25]. Likewise, cannabinoid receptors can mediate the regulation of immune function. Thus many in vitro and in vivo studies aim to exploit the putative therapeutic potential of cannabinoid signaling in inflammation-accompanied diseases (e.g., multiple sclerosis) [26]. Chemokine receptors have been identified as interesting targets for antineoplastic therapies since they are expressed on the surface of both tumor and immune cells [27]. They regulate proliferation, cell survival, differentiation, immune cell activation, cell migration, and death. Since there exist both, pro- and anti-inflammatory cytokines, they can either repress tumor growth or induce cell transformation and malignancy, always depending on the current state of the tumor microenvironment [28]. Since the immune system in general plays a critical role in various physiological and pathological processes, such as inflammation, autoimmunity, tumor growth, metastasis etc. there is a huge potential for drug discovery and development in exploiting the listed target classes further.

In the case of kinases, investigations on targets annotated with “immune system process” have also increased during the investigated period (Table 1). Notably, the main contributing class of proteins for this emerging trend are Janus kinases (JAKs) in recent years (Figure 5), here especially tyrosine kinase 2 (TYK2, JAK-A and JAK-B; Supplementary Figure S2). This class of non-receptor tyrosine kinases [29] are essential mediators for downstream signaling via STAT (signal transducer of activation) proteins and require a soluble factor to associate extracellularly with the corresponding transmembrane receptor for their activation, including cytokines and hormones. Since the JAK/STAT pathway is the major paradigm for membrane to nucleus signaling, it is also recognized as an important regulator of the immune system including the resistance to infection and maintenance of immune tolerance [30, 31]. JAKs are therefore targeted by many modern pharmaceuticals for e.g. treatment of autoimmune diseases such as myelofibrosis and rheumatoid arthritis. In addition to their role in immunity and inflammation, especially TYK2 plays an important role in the prevention and activation of carcinogenesis as well as unexpected tumor rejection [32].

**Figure 5.**
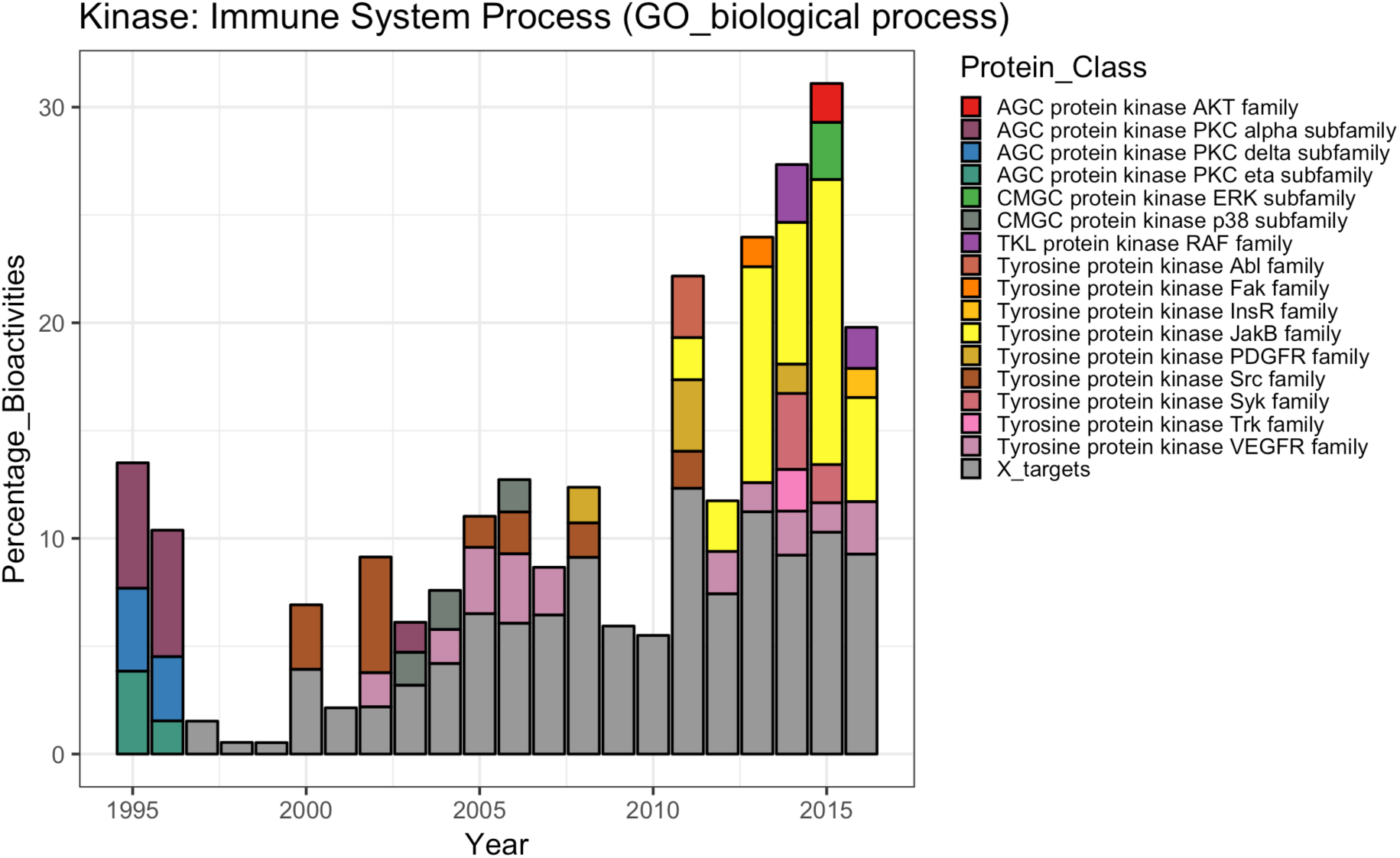
Bar chart showing the proportions of different protein classes contributing to the GO biological process annotation trend “immune system process” for kinases. Grey bars indicate target classes with little contribution to the general trend (less than 10% of the largest portion of a protein class in a certain year).

While attention in GPCR protein classes involved in “immune system processes” increased since 1995, the research interest in protein classes involved in “circulatory system processes” (definition: organ system process carried out by any of the organs or tissues of the circulatory system) decreased within the same time period for GPCR proteins. Specifically, a shrinking percentage of bioactivity data was reported for dopamine receptors, adrenergic receptors, and adenosine receptors (Figure 6). Such a trend points to decreasing interest in GPCRs as direct drug targets for cardiovascular diseases. Instead, molecular targets which are part of the signaling pathways that these GPCRs elicit (such as beta-arrestins) are nowadays investigated as promising new avenues for treating heart diseases [33].

**Figure 6.**
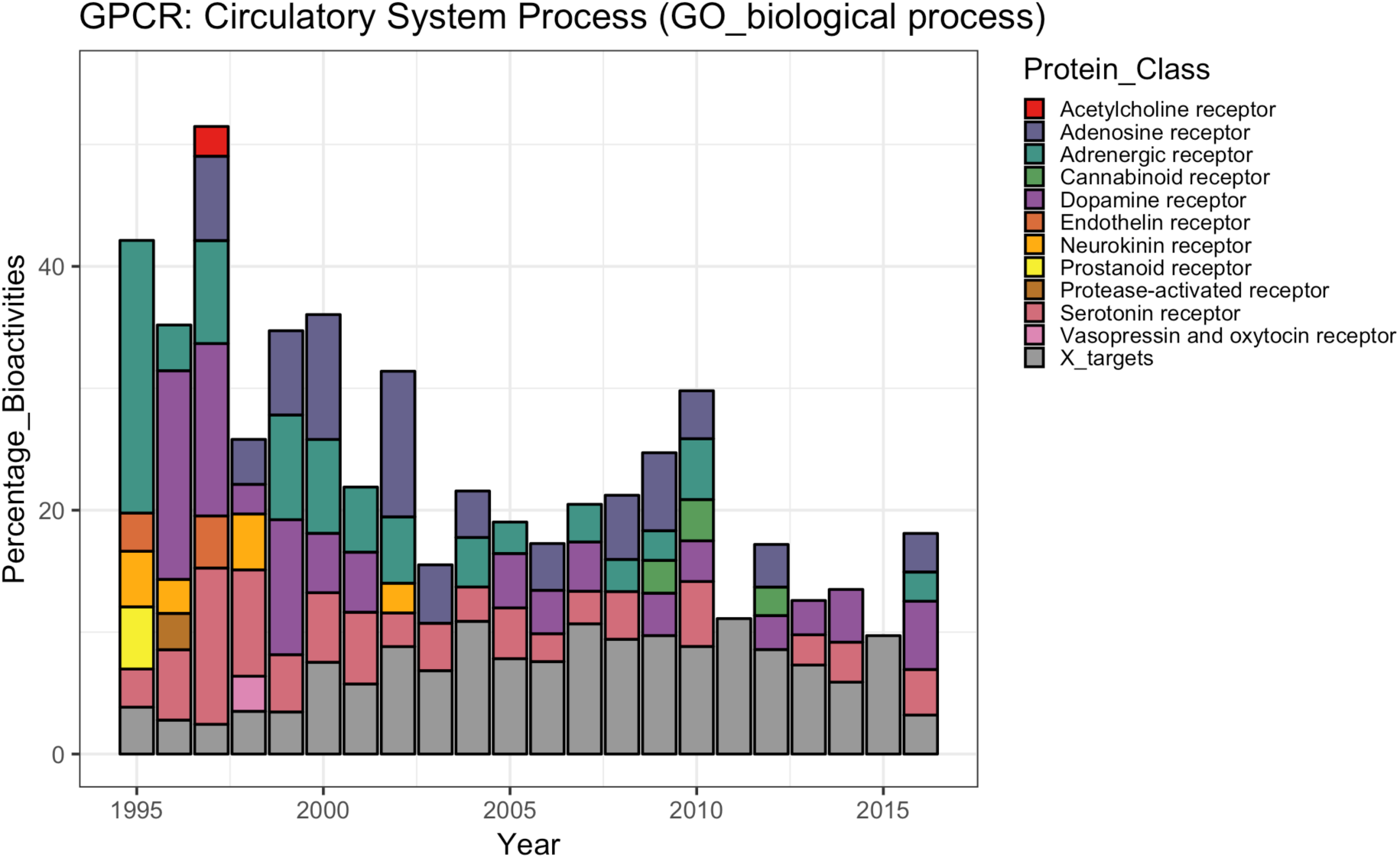
Bar chart showing the proportions of different target classes contributing to the GO biological process annotation trend “circulatory system process” for GPCR targets. Grey bars indicate target classes with little contribution to the general trend (less than 10% of the largest portion of a target class in a certain year).

Interestingly, when inspecting trends in the higher level GO term “signal transduction”, which delivered one of the steepest positive GO annotation trends for kinases and a slightly negative trend for GPCRs (see Table 1), familiar protein classes mainly contribute to these trends. For kinases, JAKs predominantly cause this steep positive trend since 2013 (Supplementary Figure S2); for GPCRs, mainly decreasing interest in adrenergic, adenosine, and dopamine receptors led to a slight decrease in bioactivity measurements on targets involved in “signal transduction” (Supplementary Figure S3).

### Evolution of disease annotations

At the level of diseases, target annotations appear to provide even more meaningful insights into target innovation trends since they delineate the extent to which the involvement of a protein (or protein class) in a certain biological process can potentially be translated into the development of pharmaceuticals for the treatment of a particular disease.

With respect to the overall distribution of the trends among the different target families, disease annotations seem to behave in analogy to GO biological process annotations: For kinases (35 diseases) and ion channels (3 diseases) exclusively positive trends were found to be statistically significant; whereas for nuclear receptors (13 diseases) and proteases (23 diseases) the disease trends are exclusively recurrent. Transporters do not follow any significant disease trends, but GPCR disease trends can be divided into positive (9 diseases) and negative (16 diseases) trends (see Table 2). No particular attempt was undertaken to filter out disease annotations that are describing tightly connected diseases (e.g. “endometrioma” and “endometriosis”) or subcategories of another disease annotation (e.g. “pancreatic neoplasm” and “malignant neoplasm of pancreas”) in order to not bias the trends by individual knowledge about certain diseases.

**Table 2.** List of annotated diseases (from DisGeNET) with statistically significant positive or negative trends (robust regression; p <= 0.05) for each target family. No significant trends could be observed for transporters.

| Target family | Disease | Slope | Target family | Disease | Slope |
| --- | --- | --- | --- | --- | --- |
| Kinase | Intellectual Disability | 0.531 | GPCR | Malignant neoplasm of pancreas | 0.177 |
| Kinase | Malignant neoplasm of breast | 0.440 | GPCR | Pancreatic Neoplasm | 0.177 |
| Kinase | Breast Carcinoma | 0.417 | GPCR | Complex Partial Status Epilepticus | 0.120 |
| Kinase | Malignant Neoplasms | 0.357 | GPCR | Grand Mal Status Epilepticus | 0.120 |
| Kinase | Schizophrenia | 0.355 | GPCR | Non-Convulsive Status Epilepticus | 0.120 |
| Kinase | melanoma | 0.353 | GPCR | Petit mal status | 0.120 |
| Kinase | Non-Small Cell Lung Carcinoma | 0.326 | GPCR | Simple Partial Status Epilepticus | 0.120 |
| Kinase | Mammary Neoplasms | 0.325 | GPCR | Status Epilepticus | 0.120 |
| Kinase | Mammary Neoplasms, Human | 0.323 | GPCR | Status Epilepticus, Subclinical | 0.120 |
| Kinase | Malignant neoplasm of prostate | 0.313 | GPCR | Congestive heart failure | -0.106 |
| Kinase | Prostatic Neoplasms | 0.312 | GPCR | Heart Decompensation | -0.106 |
| Kinase | Adenocarcinoma of lung (disorder) | 0.293 | GPCR | Heart Failure, Right-Sided | -0.106 |
| Kinase | Conventional (Clear Cell) Renal Cell Carcinoma | 0.285 | GPCR | Heart failure | -0.106 |
| Kinase | Experimental Hepatoma | 0.268 | GPCR | Left-Sided Heart Failure | -0.106 |
| Kinase | Hepatoma, Morris | 0.268 | GPCR | Myocardial Failure | -0.106 |
| Kinase | Hepatoma, Novikoff | 0.268 | GPCR | Panic Attacks | -0.127 |
| Kinase | Liver Neoplasms, Experimental | 0.268 | GPCR | Panic Disorder | -0.127 |
| Kinase | Lung Neoplasms | 0.226 | GPCR | Depression, Bipolar | -0.164 |
| Kinase | Malignant neoplasm of lung | 0.226 | GPCR | Autistic Disorder | -0.228 |
| Kinase | Malignant neoplasm of liver | 0.198 | GPCR | Intellectual Disability | -0.255 |
| Kinase | Malignant neoplasm of ovary | 0.189 | GPCR | Hypertensive disease | -0.266 |
| Kinase | ovarian neoplasm | 0.175 | GPCR | Bipolar Disorder | -0.369 |
| Kinase | Squamous cell carcinoma | 0.171 | GPCR | Bradycardia | -0.379 |
| Kinase | Liver carcinoma | 0.169 | GPCR | Hypotension | -0.467 |
| Kinase | Myocardial Ischemia | 0.153 | GPCR | Mood Disorders | -0.583 |
| Kinase | Adenocarcinoma | 0.148 | Protease | Giant Cell Glioblastoma | -0.088 |
| Kinase | Adenocarcinoma, Basal Cell | 0.148 | Protease | Glioblastoma | -0.088 |
| Kinase | Adenocarcinoma, Oxyphilic | 0.148 | Protease | Glioblastoma Multiforme | -0.088 |
| Kinase | Adenocarcinoma, Tubular | 0.148 | Protease | Thrombosis | -0.116 |
| Kinase | Carcinoma, Cribriform | 0.148 | Protease | Thrombus | -0.116 |
| Kinase | Carcinoma, Granular Cell | 0.148 | Protease | Myocardial Infarction | -0.118 |
| Kinase | Esophageal Neoplasms | 0.145 | Protease | Brain Ischemia | -0.138 |
| Kinase | Malignant neoplasm of esophagus | 0.145 | Protease | Cerebral Ischemia | -0.138 |
| Kinase | Giant Cell Glioblastoma | 0.137 | Protease | Obesity | -0.139 |
| Kinase | Neoplasm Metastasis | 0.129 | Protease | Liver carcinoma | -0.140 |
| Ion channel | Benign Neoplasm | 0.199 | Protease | Schizophrenia | -0.183 |
| Ion channel | Malignant Neoplasms | 0.199 | Protease | Colorectal Cancer | -0.200 |
| Ion channel | Neoplasms | 0.199 | Protease | Non-alcoholic Fatty Liver Disease | -0.208 |
| Nuclear receptor | Colorectal Cancer | -0.118 | Protease | Nonalcoholic Steatohepatitis | -0.208 |
| Nuclear receptor | Endometrioma | -0.137 | Protease | Neoplasm Metastasis | -0.241 |
| Nuclear receptor | Endometriosis | -0.137 | Protease | Oral Submucous Fibrosis | -0.244 |
| Nuclear receptor | Adenocarcinoma | -0.139 | Protease | Cerebral Hemorrhage | -0.258 |
| Nuclear receptor | Adenocarcinoma, Basal Cell | -0.139 | Protease | Liver Cirrhosis, Experimental | -0.322 |
| Nuclear receptor | Adenocarcinoma, Oxyphilic | -0.139 | Protease | Neoplasm Invasiveness | -0.382 |
| Nuclear receptor | Adenocarcinoma, Tubular | -0.139 | Protease | Malignant neoplasm of breast | -0.421 |
| Nuclear receptor | Carcinoma, Cribriform | -0.139 | Protease | Breast Carcinoma | -0.429 |
| Nuclear receptor | Carcinoma, Granular Cell | -0.139 | Protease | Mammary Neoplasms | -0.429 |
| Nuclear receptor | Breast Carcinoma | -0.208 | Protease | Mammary Neoplasms, Human | -0.429 |
| Nuclear receptor | Mammary Neoplasms | -0.208 |  |  |  |
| Nuclear receptor | Mammary Neoplasms, Human | -0.208 |  |  |  |
| Nuclear receptor | Malignant neoplasm of breast | -0.299 |  |  |  |

Across all target families, targets annotated to different types of cancer are by far the most frequently investigated ones as they show steep increasing trends in research attention over 22 years. This not only accounts for kinases, but also for ion channels, with a single unique disease annotation – “benign/malignant neoplasms” (Table 2 and Supplementary Figure S4).

Studying the level of individual protein classes contributing to steep bioactivity trends for cancer-related targets it becomes obvious that research was not focused on only a few protein classes. Instead, depending on cancer type diverging classes of kinases seem to be a focus of interest in recent times (Figure 7, Figure 8, and Supplementary Figure S5). However, a few protein classes are more frequently investigated across the 22 cancer related disease trends in recent years (Table 2): epidermal growth factor receptor (EGFR) class, vascular endothelial growth factor receptor (VEGFR) class, RAF kinases, and the proto-oncogene MET kinases.

**Figure 7.**
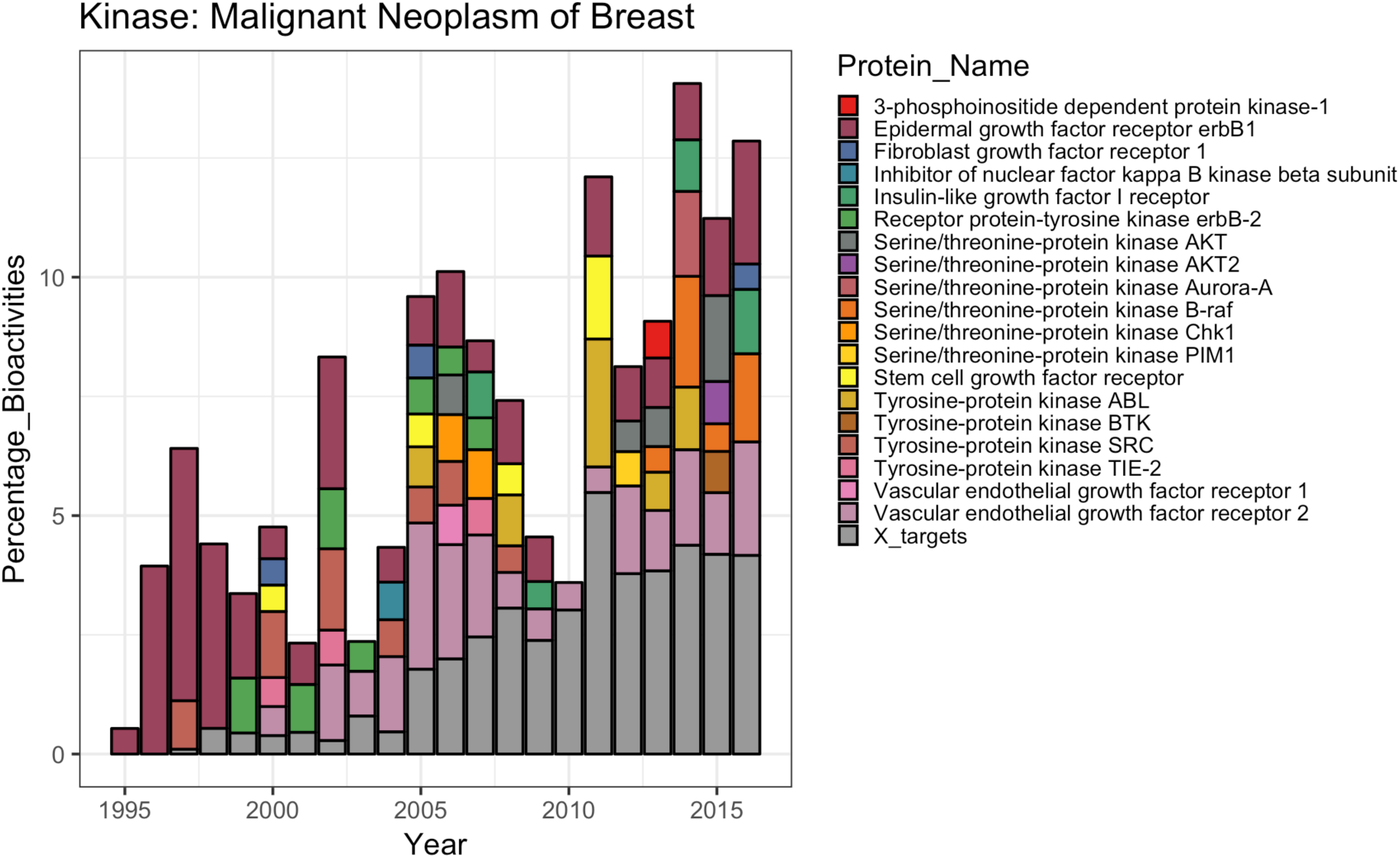
Bar chart showing the proportions of different kinases contributing to steep positive disease trend “malignant neoplasm of breast”.

**Figure 8.**
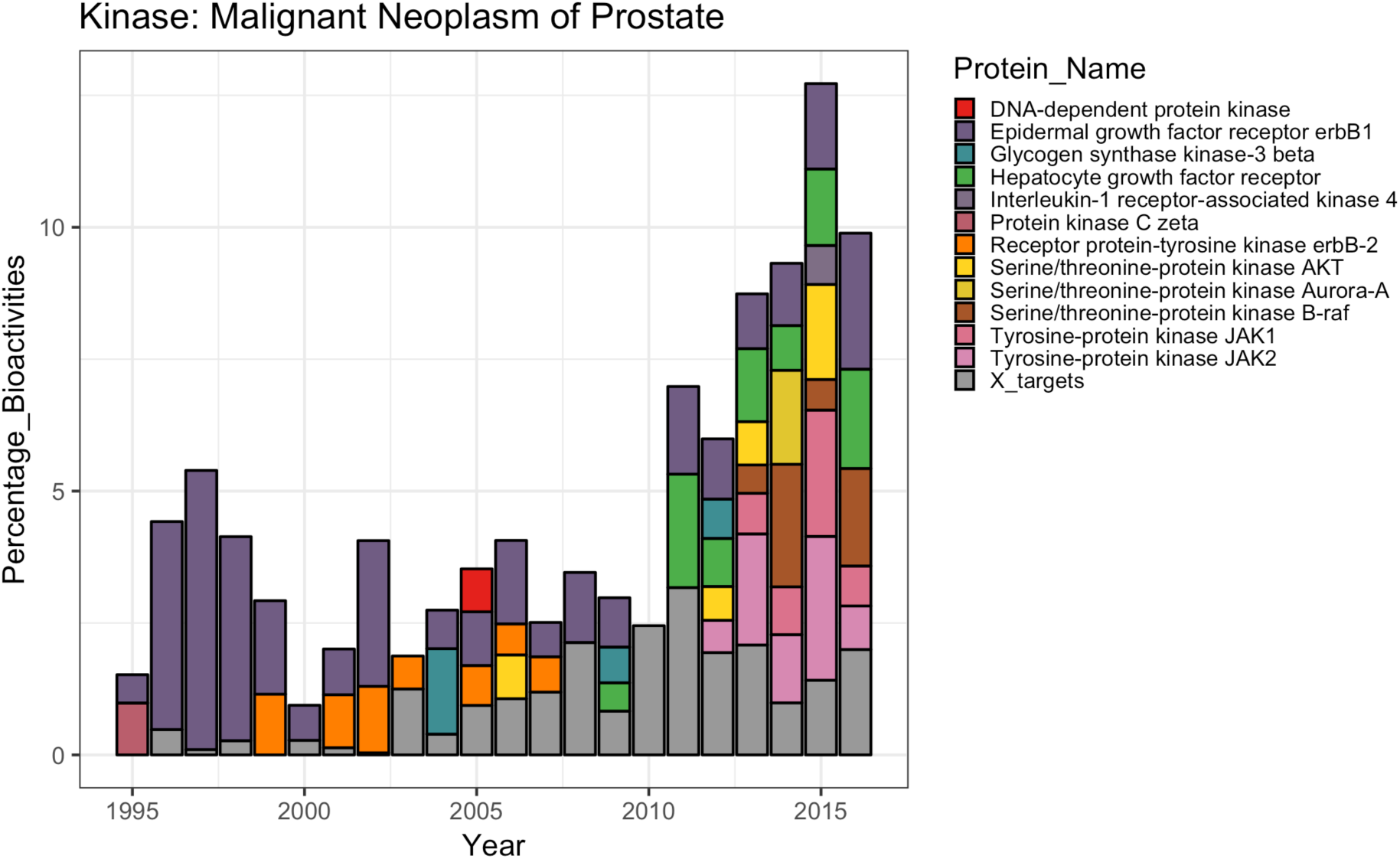
Bar chart showing the proportions of different kinases contributing to steep positive disease trend “malignant neoplasm of prostate”.

Whereas EFGR, VEGFR, and MET are receptor tyrosine kinases (RTKIs) expressed at cell-surfaces, RAF is a downstream kinase transmitting the signal to the cell nucleus. Interest in ligands targeting RAF is a more recent trend (Figure 7, Figure 8, and Supplementary Figure S5) which reflects emerging approaches in cancer research which are designed to target at the pathway level, ideally by the use of multitargeted TKIs. One prominent example is sorafenib, which inhibits VEGFR, PDGFR-(beta-type platelet-derived growth factor receptor), C-RAF, B-RAF, and c-KIT [34]. A major problem in cancer therapy is the eventual development of resistance to neoplastic therapy, which at the level of targeted anticancer therapy can be attributed to multiple resistance mechanisms including single point mutations, as well as the activation of parallel salvage pathways. Thus, a recent strategy in the development of newer antineoplastic agents is the simultaneous inhibition of multiple pathways. For example CUDC-101 – currently in phase I clinical trials [35] – synergistically blocks key regulators of EGFR/HER2 signaling pathways, as well as multiple compensatory pathways, such as AKT, HER3, and MET [36]. The increasing interest in VEGFR as target for antineoplastic drugs is attributed to its effect in preventing tumor-neovascularization (antiangiogenic therapy) [37].

The JAKB family which has been identified as the major contributor to the steep GO biological process trend “immune system process” for kinases only appeared to contribute to the emerging trend in the case of “malignant neoplasm of prostate”, “prostatic neoplasms”, and “experimental hepatoma” (Figure 8 and Supplementary Figure S5). At the level of the individual targets it becomes clear that for research on targets annotated to the disease e.g. “malignant neoplasm of prostate” the epidermal growth factor receptor ErbB1 used to be in the main focus of interest until approx. 2009. Later, other protein targets such as the Hepatocyte growth factor receptor, B-RAF, as well as the Janus kinases JAK1 and JAK2 received more and more interest (Figure 8).

Besides cancer-related diseases, “intellectual disability” (subnormal intellectual functioning which originates during the developmental period; previously referred to as “mental retardation”) [38] and “schizophrenia” are also among the steep positive disease trends for kinases (Table 2, Supplementary Figure S6 and S7). Interestingly, for one of the main protein targets contributing to both positive disease trends in recent years – the hepatocyte growth factor receptor (c-MET) – associations to the neurodevelopmental disease “intellectual disability” are not described in the scientific literature to date, but were retrieved merely from the publicly available knowledge base “Genomics England PanelApp” [39] (via DisGeNET) [40]. The same accounts for the association between JAK3 and “intellectual disability”. Such findings are especially interesting in the context of novel therapeutic opportunities and drug repurposing since a strong genetic link between the target and the disease is confirmed but research in the respective indication area is still in its childhood. For schizophrenia, however, an influence of mutations in the MET proto-oncogen was investigated systematically since a lower incidence of cancer in schizophrenia patients was observed previously [41]. The authors also postulate an influence of MET variation in neurocognition and neurodevelopmental processes in general, since MET was also found to be involved in autism spectrum disorder (ASD) [42]. In contrast, other protein-disease associations contributing to this steep trends for “intellectual disability” and “schizophrenia” are better documented in the scientific literature to date, such as the involvement of the serine/threonine kinase AKT in these diseases [43, 44]. Vascular endothelial growth factor receptor 2 (VEGFR2) is the most heavily studied protein that is associated with “schizophrenia”. The growth factor binding to this receptor (VEGFA) has been implicated in neurotrophy and neurogenesis as well in the pathophysiology of schizophrenia [45].

In the case of ion channels, the class of voltage-gated potassium channels is mainly contributing to the increasing disease trend “neoplasms” (identical to the trends for “malignant neoplasms” and “benign neoplasm”), and to a lesser extent transient receptor potential (TRP) channels. Interestingly, there is only a single representative from each class contributing to this trend, respectively: HERG and the vanilloid receptor (Supplementary Figure S9). Over- and/or mis-expression of HERG in tumor cells has been identified as a critical factor for establishment and maintenance of neoplastic growth [46]. Thus, HERG inhibitors can reduce proliferation both in vitro and in vivo which offers new antineoplastic strategies [47]. Also, vanilloid receptors have been found to regulate cancer cell proliferation, angiogenesis and apoptosis and thus have been suggested as potential anticancer targets and prognostic markers in recent years [48].

For GPCRs, just two diseases show a positive trend: pancreatic neoplasm (identical trend to “malignant neoplasm of pancreas”) and different forms of epilepsy (all forms are describing identical trends). Notably, for nuclear receptors and proteases cancer-related diseases are showing a recurrent trend (Table 2 and Supplementary Figure S5). Cancer research thereafter tends to shift to mainly kinase targets while research at nuclear receptor and protease targets in the domain of cancer likely diminished over the past 22 years.

As already apparent from GO biological process annotations, interest in GPCR targets associated to cardiovascular diseases decreased which is confirmed by the declining interest in GPCRs associated with cardiovascular diseases (such as “hypotension”, “bradycardia”, and “hypertensive disease”). In addition, mood disorders seem to be subjected to declining attention in GPCR research (for individual target contributions to steep positive and negative disease trends for GPCRs see Supplementary Figure S10).

### Dynamic networks to study the evolution of associations between targets, GO terms, and disease annotations – Uncovering new links between biological processes and diseases

In the previous sections, we have described 1) target families of emerging interest in drug discovery (by tracking target innovation trends comparing different protein families), 2) biological processes of increasing attention (in the form of GO annotations), as well as 3) trends of annotated diseases per target family. Next, through linkage of targets, GO terms, and diseases, the otherwise disparate knowledge in the domain of small molecule drug discovery is pooled in order to unravel new potential research directions. Since GO biological process annotations and disease annotations were in first instance mapped independently via protein targets, such unbiased exploration of network trends can deliver insights into biological processes that are associated with a particular disease. Vice versa, by filtering for a particular GO annotation, diseases and related targets can be explored.

In order to include the dimension of time like in the precedent investigations, the study period was split into two equal parts (11 years each) and consequently two independent networks were generated for each investigated disease or GO annotation. Thus, in conjunction, the network views provide a “dynamic network” perception which delineates major changes in network connectivity over time.

Figures 9-11 are showing the protein class-GO annotation networks for two selected emerging diseases from the kinase data set: “malignant neoplasm of breast”, and “malignant neoplasm of prostate”. A study of the networks for the two cancer types (Figure 9 and 10) reveals important shifts in the foci of investigated targets over time. For both cancer-type malignancies, proteins belonging to the class of epidermal growth factor receptors (short name “Egfr” in graphics) have been the most heavily investigated targets (in terms of percentages of bioactivities) in the earlier period (from 1995-2005). In the later period (2006-2016), other protein classes outpaced the class of EGF-receptors (although EGFRs are still being ranked second): in case of the disease “malignant neoplasm of breast” the class of vascular endothelial growth factor receptors (short name “Vegfr” in graphics), in the case of “malignant neoplasm of prostate” the class of Janus kinases are listed as most heavily investigated target classes with an annotation to the respective disease. Supplementary Tables 2 - 5 list all target classes associated with these diseases and their absolute and relative amounts of bioactivities in the respective time periods.

**Figure 9:**
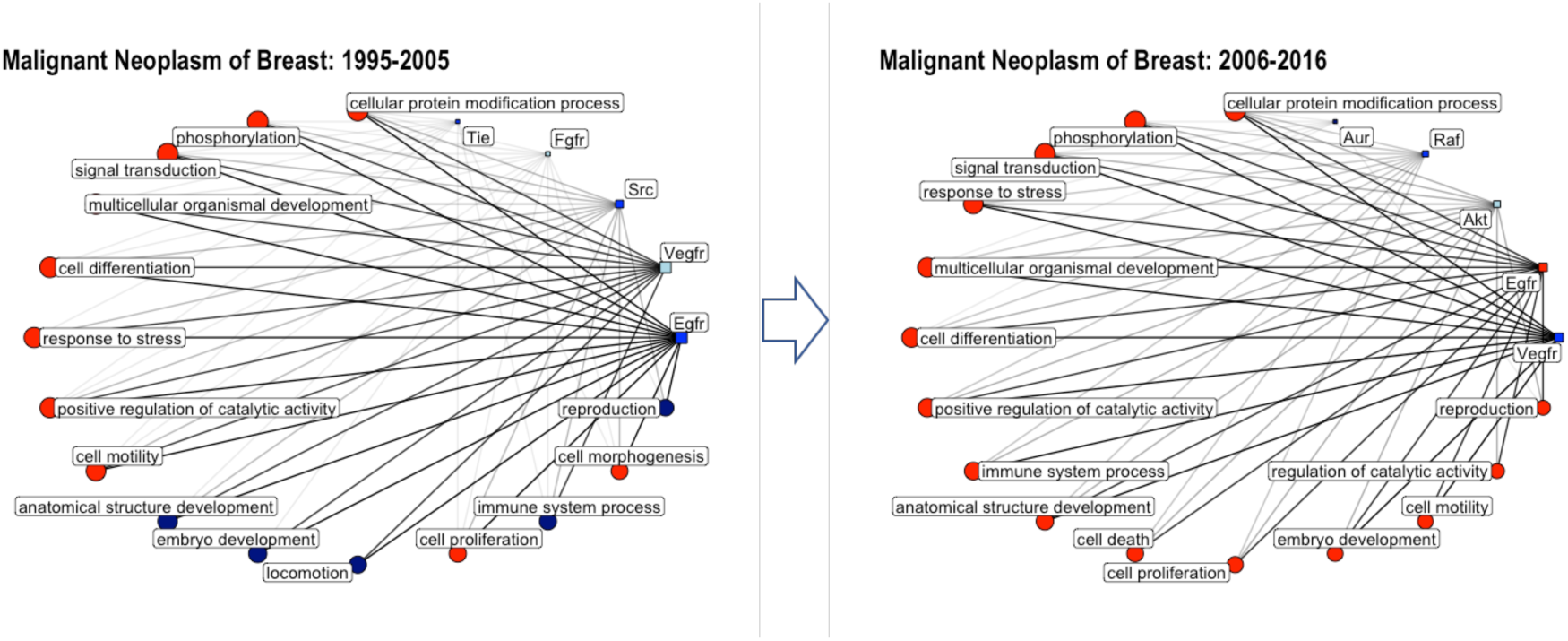
Network graphs showing the connectivity of kinases and GO biological process annotations annotated to the disease “malignant neoplasm of breast” for the time periods 1995-2005 and 2006-2016. Circles correspond to GO biological process annotations; squares correspond to protein classes; only the top 5 target classes and top 15 GO terms are shown (with respect to percentage of total bioactivities); the size of the circle/square as well as the thickness of the connecting lines is correlated to the percentage of bioactivities associated with this GO annotation/protein class; the color code is reflecting the number of available drug-efficacy target annotations (light blue: 0; blue: 1-5; navy: 6-10; red >10); Akt: AGC protein kinase AKT family; Aur: Other protein kinase AUR family; Egfr: Tyrosine protein kinase EGFR family; Fgfr: Tyrosine protein kinase FGFR family; Raf: TKL protein kinase RAF family; Src: Tyrosine protein kinase Src family; Tie: Tyrosine protein kinase Tie family; Vegfr: Tyrosine protein kinase VEGFR family.

Interestingly, when plotting the network with single protein targets (instead of protein classes; Supplementary Figures S10 and S11), this trend is less pronounced since ErbB1 as a main contributor from the EGFR family keeps being listed second for “malignant neoplasm of breast” and first for “malignant neoplasm of prostate”. Thus, obviously only interest in ErbB2 decreased over time (disappearing from the list of top 10 ranked proteins in case of “malignant neoplasm of breast”; moving from rank position 2 to position 9 for “malignant neoplasm of prostate”). These trends were already visible from the stacked bar plots showing the target composition of these diseases over time (Figures 7 and 8). However, in addition to this information, dynamic network representations do also provide insights into the biological processes that the emerging targets for a particular disease are involved in and in particular they are showing potential changes in the prevalence of the GO annotations over time. Such networks can provide a focused view on the interconnectivity of targets and biological processes for a particular disease and they facilitate the comparison of the evolution of these relationships over time. For instance, in the case of the cancer-related diseases chosen as examples here (Figures 9 and 10) we can observe an increasing interest in targets involved in “immune system processes”. For targets annotated to “malignant neoplasm of breast”, mainly the protein classes VEGFR, AKT, and RAF (single targets: VEGFR2, SRC, ABL, Insulin-like growth factor receptor I, B-RAF, and AKT; see Figure 7 and Supplementary Figure S10) are associated to this GO term within the period 2006-2016. For targets annotated to “malignant neoplasm of prostate”, JakB, AKT, and RAF are the predominant contributors (single targets: JAK2, AKT, B-RAF, Aurora-A; see Figure 8 and Supplementary Figure S11).

**Figure 10:**
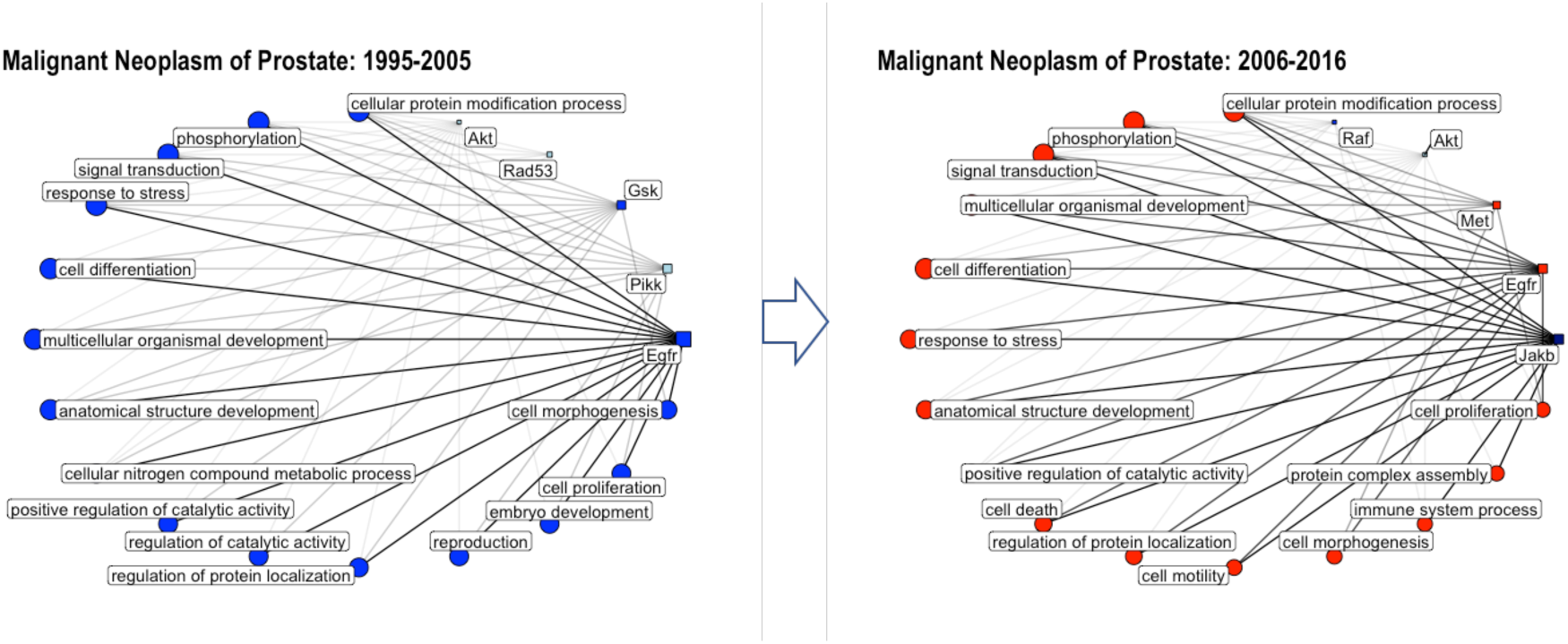
Network graphs showing the connectivity of kinases and GO biological process annotations annotated to the disease “malignant neoplasm of prostate” for the time periods 1995-2005 and 2006-2016. Circles correspond to GO biological process annotations; squares correspond to protein classes; only the top 5 target classes and top 15 GO terms are shown (with respect to percentage of total bioactivities); the size of the circle/square as well as the thickness of the connecting lines is correlated to the percentage of bioactivities associated with this GO annotation/protein class; the color code is reflecting the number of available drug-efficacy target annotations (light blue: 0; blue: 1-5; navy: 6-10; red >10). Akt: AGC protein kinase AKT family; Egfr: Tyrosine protein kinase EGFR family; Gsk: CMGC protein kinase GSK family; Jakb: Tyrosine protein kinase JakB family; Met: Tyrosine protein kinase Met family; Pikk: Atypical protein kinase PIKK family; Rad53: CAMK protein kinase RAD53 family; Raf: TKL protein kinase RAF family.

This agglomerated knowledge on targets being involved in a particular disease and leveraging (a) particular biological process(es) can be used for designing multi-targeted drugs with an enhanced modulatory effect. Such visualizations can therefore provide an orthogonal level of systems biology insights to a view that includes pathway-information (i.e. on up- and downstream proteins). Since the number of available drug-efficacy target annotations is given by a color code, protein targets of emerging interest for a particular disease but with a small number of available drugs can be identified easily. Such targets might offer new treatment options, e.g. Serine/threonine kinase AKT for different cancer types [49] (see Figures 9 and Figure 10; Supplementary Figure S10 and S11).

In contrast, dynamic protein class-disease networks for a particular biological process (GO term) over time as exemplified in Figure 11 for “immune system process” delivers a comprehensible overview of the degree of interest in target classes that might be leveraged for operating through the particular process. By creating a single network for multiple target families (for kinases and GPCRs in Figure 11) dominant targets/target classes across families can be identified, such as e.g. cannabinoid receptors and Janus kinases in the more recent period for “immune system processes” (Figure 11). A list of all target classes annotated to “immune system process”, including their absolute and relative bioactivity amounts within the two time periods, is given in Supplementary Tables S6 and S7. Although appearing in the same network, cannabinoid receptors and Janus kinases are showing differences in their associations to diseases. Cannabinoid receptors are linked to schizophrenia, different forms of breast cancer, and unipolar depression (to name just the top ranked diseases). In contrast, Janus kinases are not linked to breast cancer, but to intellectual disability (which in return is not linked to cannabinoid receptors). The association between Janus kinases and intellectual disability is a quite recent finding since the role of the JAK-STAT pathway in neuronal differentiation and gliogenesis is only insufficiently understood to date [50].

Looking for possible links between diseases at the level of a conjoint biological process (“immune system process”), it becomes obvious that the cannabinoid receptor CB2 and VEGFR2 are both strongly linked to breast cancer as well as schizophrenia (Supplementary Figure S12). A link between cancer and schizophrenia has been proposed earlier [41, 51], as well as an influence of CB2 activation or inhibition on VEGFR2-signalling [52, 53]. The identical target-connectivity of these classification-wise disparate pathologies suggests similar treatment options for both. While the use of the antipsychotic drug pimozide for the treatment of schizophrenia is under active investigation for more than two decades [54], it was only recently reported as a potential new route for treating triple-negative breast cancer [55]. Pimozide suppressed cell growth of breast cancer cells in vitro and blockage of VEGFR2 was suggested to be responsible for inhibiting tumor development [55].

**Figure 11:**
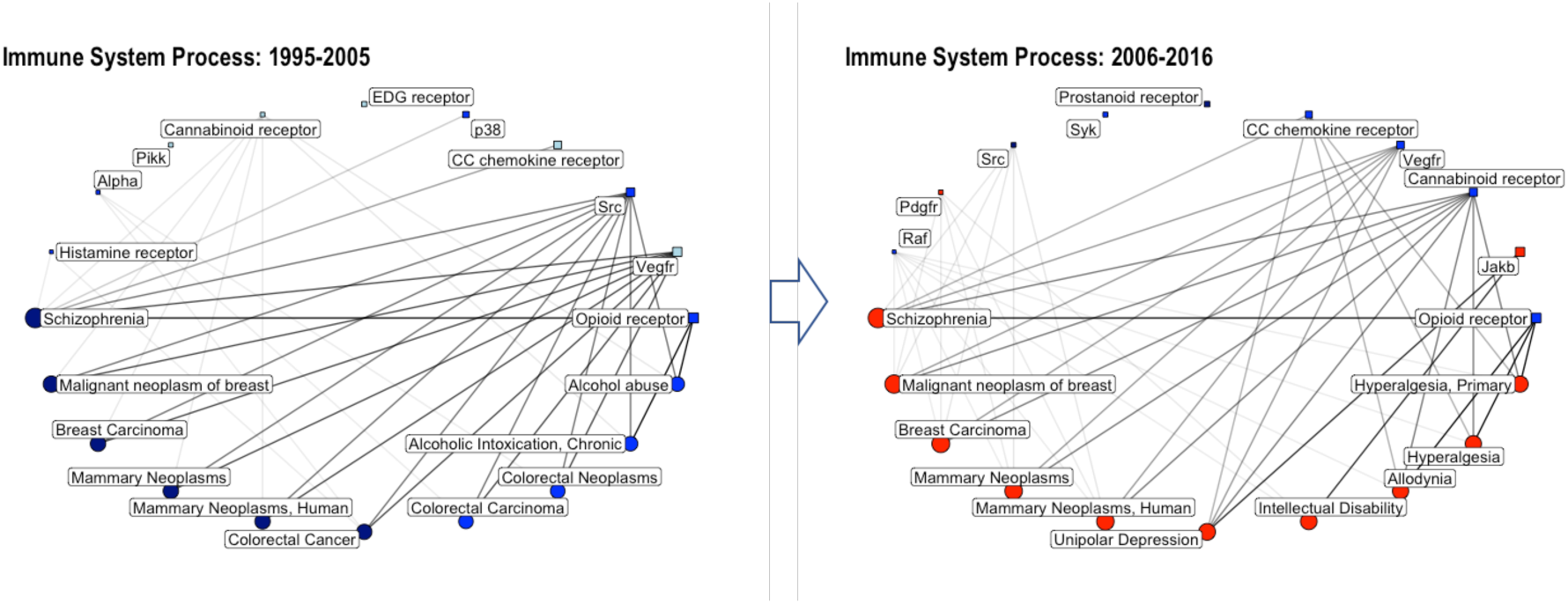
Network graph showing the connectivity of GPCRs/kinases and diseases annotated to the GO biological process “immune system process” for the time periods (from left to right): 1995-2005 and 2006-2016. Circles correspond to disease annotations; squares correspond to protein classes; only the top 10 target classes (from the merged kinase-GPCR data set) and top 10 disease annotations are shown (with respect to percentage of total bioactivities); the size of the circle/square as well as the thickness of the connecting lines is correlated to the percentage of bioactivities associated with this disease/protein class; the color code is reflecting the number of available drug-efficacy target annotations (light blue: 0; blue: 1-5; navy: 6-10; red >10). Alpha: AGC protein kinase PKC alpha subfamily; Jakb: Tyrosine protein kinase JakB family; p38: CMGC protein kinase p38 subfamily; Pdgfr: Tyrosine protein kinase PDGFR family; Pikk: Atypical protein kinase PIKK family; Raf: TKL protein kinase RAF family; Src: Tyrosine protein kinase Src family; Syk: Tyrosine protein kinase Syk family; Vegfr: Tyrosine protein kinase VEGFR family.

**Figure 12:**
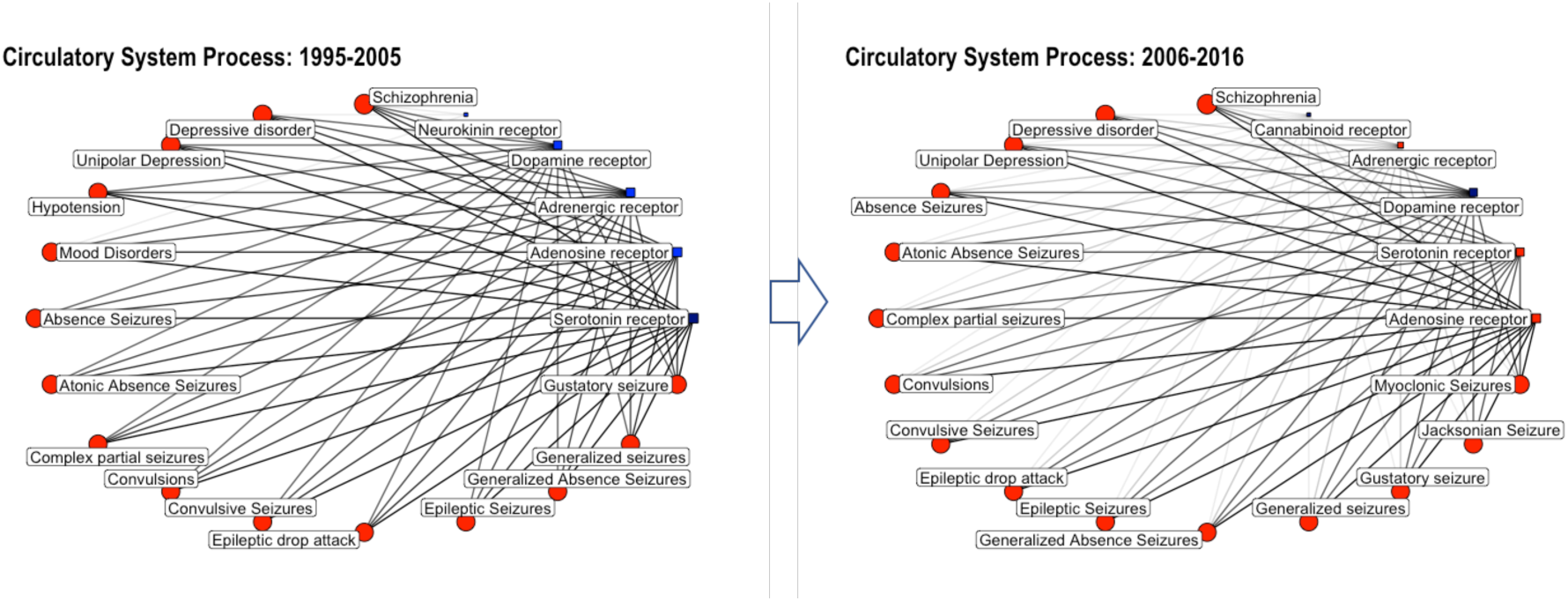
Network graph showing the connectivity of GPCRs and diseases annotated to the GO biological process “circulatory system process” for the time periods (from left to right): 1995-2005 and 2006-2016. Circles correspond to disease annotations; squares correspond to protein classes; only the top 5 target classes and top 15 disease annotations are shown (with respect to percentage of total bioactivities); the size of the circle/square as well as the thickness of the connecting lines is correlated to the percentage of bioactivities associated with this disease/protein class; the color code is reflecting the number of available drug-efficacy target annotations (light blue: 0; blue: 1-5; navy: 6-10; red >10).

Inspecting target trends of target classes annotated to “circulatory system processes” confirms the previous observation of declining research interest of GPCR proteins associated to cardiovascular diseases: e.g., target classes and single protein targets annotated to hypotension show declining interest nowadays (visible from Figure 11 and Supplementary Figure S13 and quantified in Supplementary Tables S8 and S9). Instead we can observe more interest in targets annotated to different forms of epilepsy (as identified earlier when inspecting the evolution of disease annotations; Table 2). Strikingly, cannabinoid CB1 receptors have caught up with dopamine D2 and adenosine A2a receptors, as visible from the single target-disease networks in Supplementary Figure S13.

The outlined examples demonstrate the potential of the method which uses data mapping and data visualization for the identification of malignancies and potential candidates for drug repurposing.

## Summary and Conclusions

In this study, different views of target innovation are discussed by studying time trends in research attention to particular targets in a protein family-wise fashion. Our study of emerging target families (in terms of numbers of bioactivities, targets, annotated drugs, and papers) highlights kinases and GPCRs as the most heavily investigated protein families. Interestingly, kinases have outpaced GPCRs in recent years which is especially pronounced in terms of investigated compounds and targets. A temporal view of the evolution of target developmental level (TDL) categories reveals areas of interest in target space that have just emerged in the scientific literature, such as T_bio_ and T_dark_ targets, where no active molecules are known. Many targets within these categories are kinases and transporters, highlighting novel opportunities for drug intervention.

Two orthogonal views of target innovation are considered by investigating trends in GO biological process terms and disease annotations over time. We note that no proof can be provided whether there is a causal link between attention being paid to a target or target family and a detected emerging disease or biological process trend. We investigated how intense a particular target was investigated and reported in the medicinal chemistry literature without extracting information on potential indication areas from the respective literature sources. Thus, the emerging biological process and disease trends rather highlight opportunities for therapeutic intervention rather than a real current focus on a particular disease or biological process.

Through data mapping in conjunction with data visualization, trends in research attention can be captured at an early stage also for researchers who are not experts in a particular research domain. In addition, not all links between targets and diseases are described in the scientific literature, but are only available from different knowledge bases and are made visible through the data mapping procedure (such as the link between “intellectual disability” and c-MET).

Importantly, the described methodology to analyse and visualize the data can only depict information that is already available. Its strength lies in the interconnection of the disparate scientific sub-domains: drug discovery and biology linked by bioinformatics analysis. This implies that the method works well when considering data-rich domains, as underrepresented targets families will tend to be ignored if one considers only the number of measurements. However, it can in principle also be applied to only a specific target family or target class in order to identify emerging trends within that particular domain.

One limitation is the availability of GO and disease annotations for particular targets which might be unevenly distributed, biasing the amount of available annotations to already heavily studied targets. An extreme example are orphan targets, where no potent compounds are known and there are few if any diseases or biological process associations.

The question if such an analysis rather contributes to a reinforcement in the study of already heavily investigated targets and diseases - rather than inspiring research on novel targets and in new therapeutic areas - remains. Most likely, the relationships between compounds, biological processes and diseases that are being unearthed by the network representations, and more specifically the dynamic changes of these relationships over time, will deliver the most useful information at the intermediate stage of developmental level of a target. For those cases, a link between the target and a disease has already been demonstrated by the available information, but drug discovery for that particular target or for the envisaged therapeutic indication should ideally be in its childhood still.

Thus - if used in a comprehensive manner - analysing the temporal “movement of targets” from a data science perspective enables researchers to look back on their way forward when planning and directing their future research.

## Methods

### Data retrieval

Data was retrieved from ChEMBL23 [56] via a PostgreSQL interface (pgAdmin3) by querying for compounds along with their bioactivity information (activity type, activity standard value, standard unit etc.), structural information (canonical smiles, standard InChI, and InChIKeys), target information (including protein classification), assay information and document information. In this initial query, filters for the target organism (“Homo sapiens”), target type (“SINGLE PROTEIN”), assay confidence score (>8), document year (>1985), and standard units for activities (“NOT NULL”) were set. This led to an initial dataset with 977,500 data entries (314,406 unique compounds).

Next, the data file was imported into a KNIME (version 3.5) [57] environment to perform data filtering, processing and analysis (details are provided in the following subsections).

### Data filtering

The raw data (ca. 978 K data points) was filtered for compounds with a MW <= 800, publication years 1995-2016, and data points with a pChEMBL value only (nanomolar unit and “=” as relation sign for bioactivity measurement). This led to a final data set of 514,110 data points (248,822 unique compounds).

### Retrieving and mapping of gene ontology annotations

A list of all protein targets with their respective protein classification as well as GO annotations was retrieved in a separate sql query (data set with 153,000 data points). Subsequent mapping of target ID’s (via the ChEMBL target tid’s) to protein classes and GO annotations was performed in KNIME. GO annotations were filtered for “GO biological process” annotations only.

### Filtering for individual protein families

The data set containing mappings for targets, GO terms, and diseases was split into six major protein classes by filtering the column “protein_class_desc” (protein class description) for *kinase*, *7tm1*, *transcription factor*, *protease*, *ion channel*, *transporter* (to filter for kinases, GPCRs, nuclear receptors (NR), proteases, ion channels, and transporter, respectively). Mapping these six subsets with the large data set (ca. 514 K data points) via tid’s led to the final data sets for the six target classes: approximately 99 K data points for kinases, 144 K for GPCRs, 28 K for NR, 42 K for proteases, 24 K for ion channels, and 20 K for transporter. In a separate step, tid - UNIPROT ID mappings were retrieved in a sql query and UNIPROT IDs were added to the final six target class data sets.

### Retrieving drug-efficacy target annotations

ChEMBL provides an annotation, called DISEASE_EFFICACY, which flags whether the target assigned is believed to play a role in the efficacy of the drug in the indication(s) for which it is approved (1 = yes, 0 = no). These annotations were retrieved from ChEMBL23 via a separate sql query and mapped via target tid’s and drug molregno’s.

### Mapping target developmental level (TDL) categories

Mappings of ChEMBL protein targets to TDL categories were kindly provided by Vishal Shiramshetty (NCATS/NIH).

### Retrieving and mapping of disease annotations

Curated gene-disease associations (81,746) and mappings of UNIPROT IDs and gene IDs were downloaded in csv format from DisGeNET and imported in KNIME. These gene-disease annotations were collected from UNIPROT, CGI, ClinGen, Genomics England, CTD (human subset), PsyGeNET, and Orphanet and provide a score to rank these associations by taking into account the number and type of sources (level of curation, organisms), and the number of publications supporting the association. Only associations with a score equal or greater than 0.3 were retained (at least 1 curated source reporting the gene-disease association; 81,386 associations). First, UNIPROT IDs were mapped to diseases via the associated gene IDs in KNIME. Then, UNIPROT IDs served to map diseases to the six target class data sets.

### Trend analyses

Data processing for subsequent trend calculation was performed in KNIME. Target innovation trends, GO term trends, and disease trends were described by a robust linear regression using time (Year) as an explanatory variable and the numbers of published bioactivities (measurements) per year as the response variable. The number of bioactivities was always related to the total number of bioactivities in the respective year, expressing it as a percentage of (total) bioactivities. Trend analyses were performed in R version 3.5.2 using the rlm function (“MASS” package version 7.3-51.1). Statistical significance of the trends was assessed by conducting a Wald test (“sfsmisc” package version 1.1-3) on the fitted model and extracting the p-value. Regression fits with p-values < 0.05 were considered significant, otherwise insignificant.

### Linking GO annotations and diseases

The separate data sets with the mapped GO terms and mapped disease annotations were linked by first filtering for a particular disease (e.g. “malignant neoplasm of breast”) or a particular GO annotation (e.g. “immune system process”) and then mapped via targets. To study the target connectivities of GO terms and diseases that have been identified to cause steep increasing or decreasing trends in research attention considering a particular target family, network representations were produced usually for data from that particular target family only (in case of “immune system process” a single network was created for data from both kinases and GPCRs). Static networks were created for two different time periods: 1995-2005 and 2006-2016. Taken together the two networks form a dynamic network representation, being able to compare changes in connectivities over time.

### Data visualization

Line plots and bar charts were produced in R by using the package “ggplot2” (version 3.1.0), network graphs by using the package “ggraph” (version 1.0.2).

The data for creating the stacked bar charts showing the proportions of different target classes contributing to a particular GO biological process or disease was prepared in KNIME beforehand. Only protein classes are shown as separate blocks if their percentage of bioactivities in a particular year is at least 10% of the largest contributing protein class (in any year). Target classes with minor contribution to a trend are denoted as “X_targets” in the visuals.

Bipartite network representations for selected diseases and GO terms were constructed by using the package “ggraph” (version 1.0.2) in R by displaying the connectivities of protein classes (=first set of nodes) and GO terms or diseases (=second set of nodes). Only the top 5 protein classes and top 15 GO terms/diseases are displayed respectively (with respect to percentages of annotated bioactivities). The disease-target class network for “immune system process” displays the top 10 protein classes, since these include both proteins from the kinase and GPCR families.

### Data, code, and workflow availability

All data sets, R code, as well as KNIME workflows used for performing the analyses are publicly available: https://github.com/BZdrazil/Moving_Targets.

## Authors’ contributions

BZ designed the study, performed the analysis, and wrote the manuscript. LR helped in designing the study and performing the analysis. NB and RG advised during the process of designing and performing the study, and helped writing the manuscript. All authors read and approved the final manuscript.

## Acknowledgements

We acknowledge NCATS/NIH for providing us with mappings of targets to their Target Developmental Level (TDL) categories.

## Supplementary Information

**Supplementary Figure S1:**
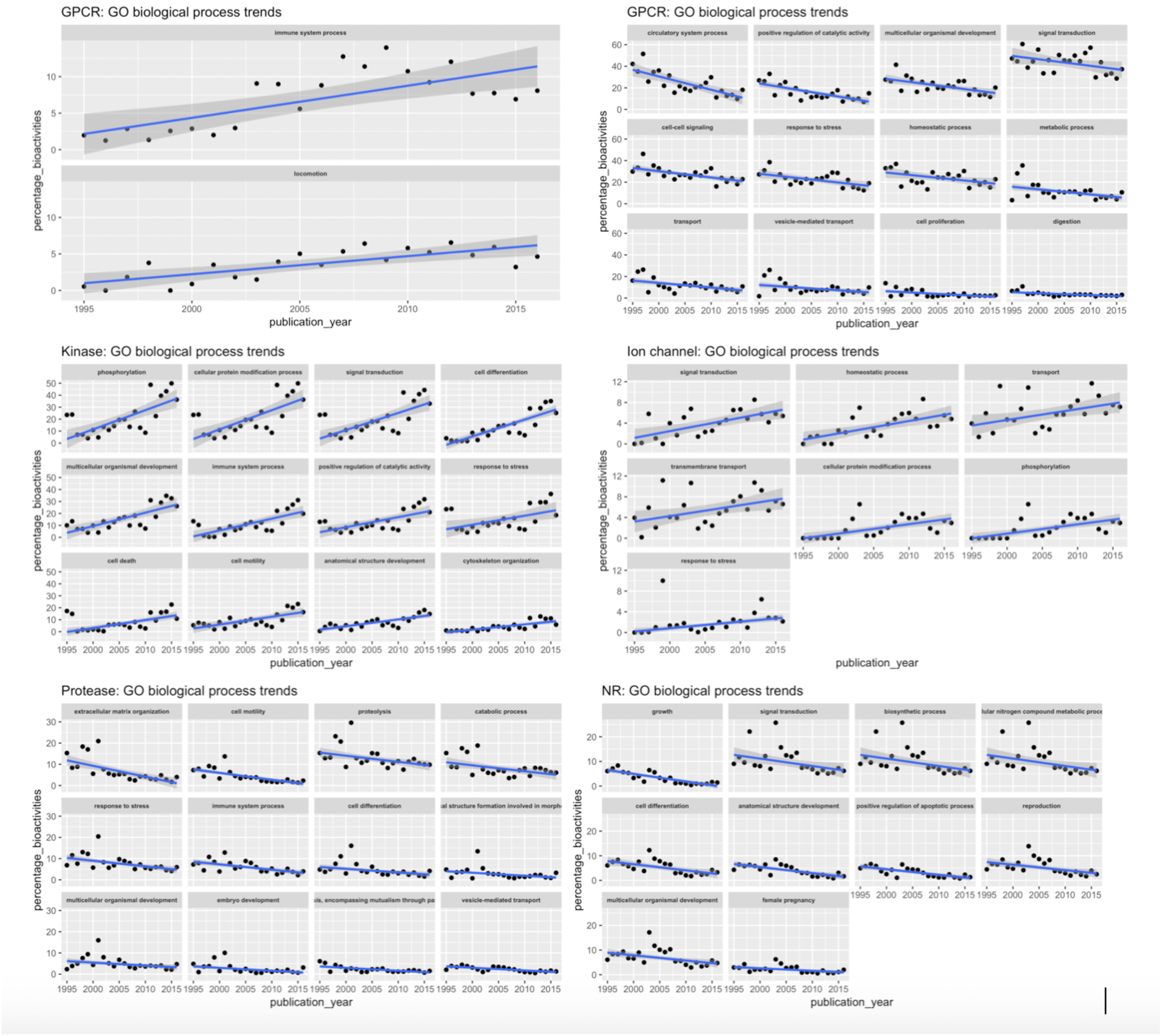
Trend line plots for GO biological process annotations (per target family). Only the twelve steepest positive and twelve steepest negative trends are shown (if available) considering only statistically significant trends (robust regression; p <= 0.05).

**Supplementary Figure S2:**
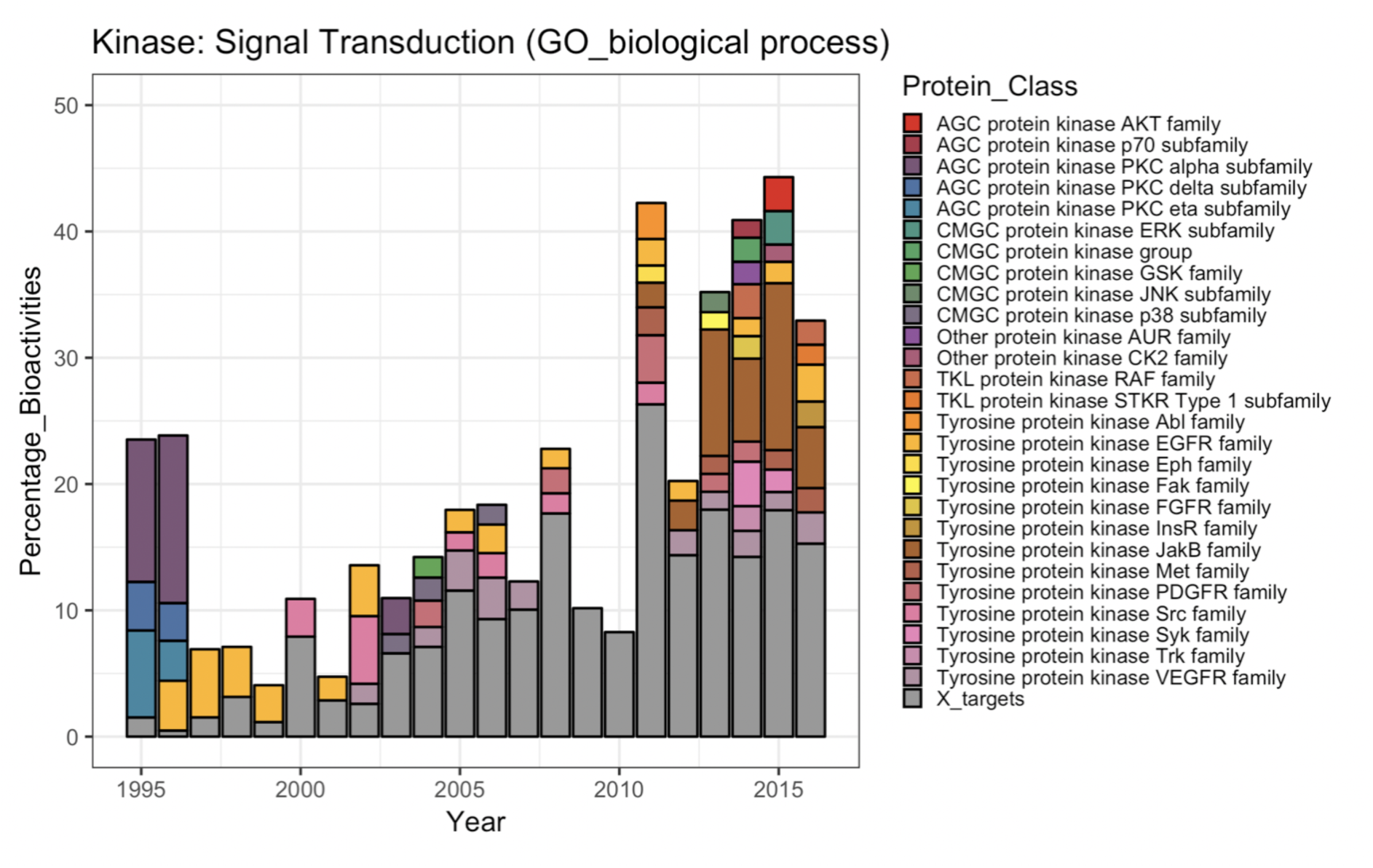
Bar chart showing the proportions of different target classes contributing to the GO biological process annotation trend “signal transduction” for kinase targets. Grey bars indicate target classes with little contribution to the general trend (less than 10% of the largest portion of a target class in a certain year).

**Supplementary Figure S3:**
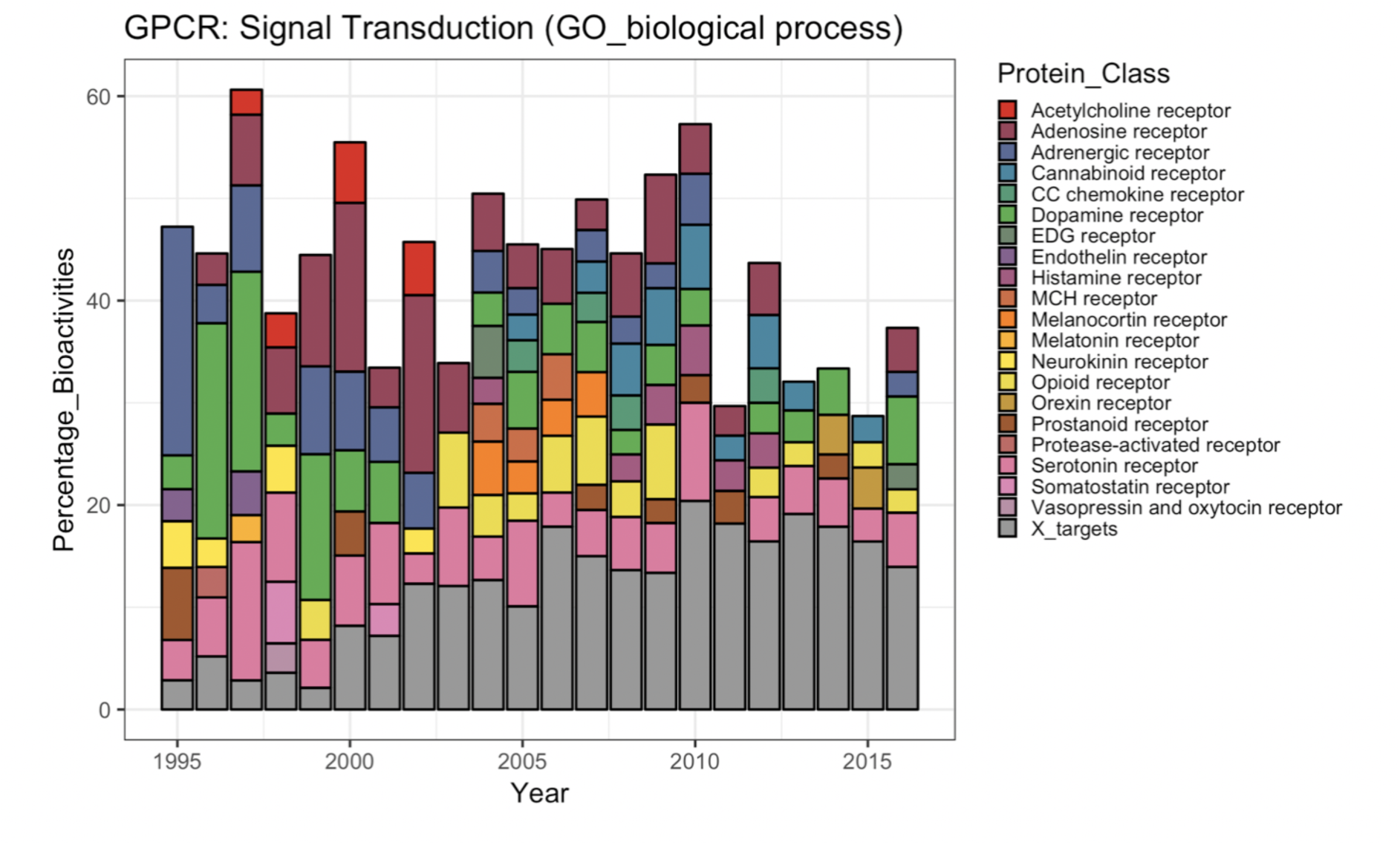
Bar chart showing the proportions of different target classes contributing to the GO biological process annotation trend “signal transduction” for GPCR targets. Grey bars indicate target classes with little contribution to the general trend (less than 10% of the largest portion of a target class in a certain year).

**Supplementary Figure S4:**
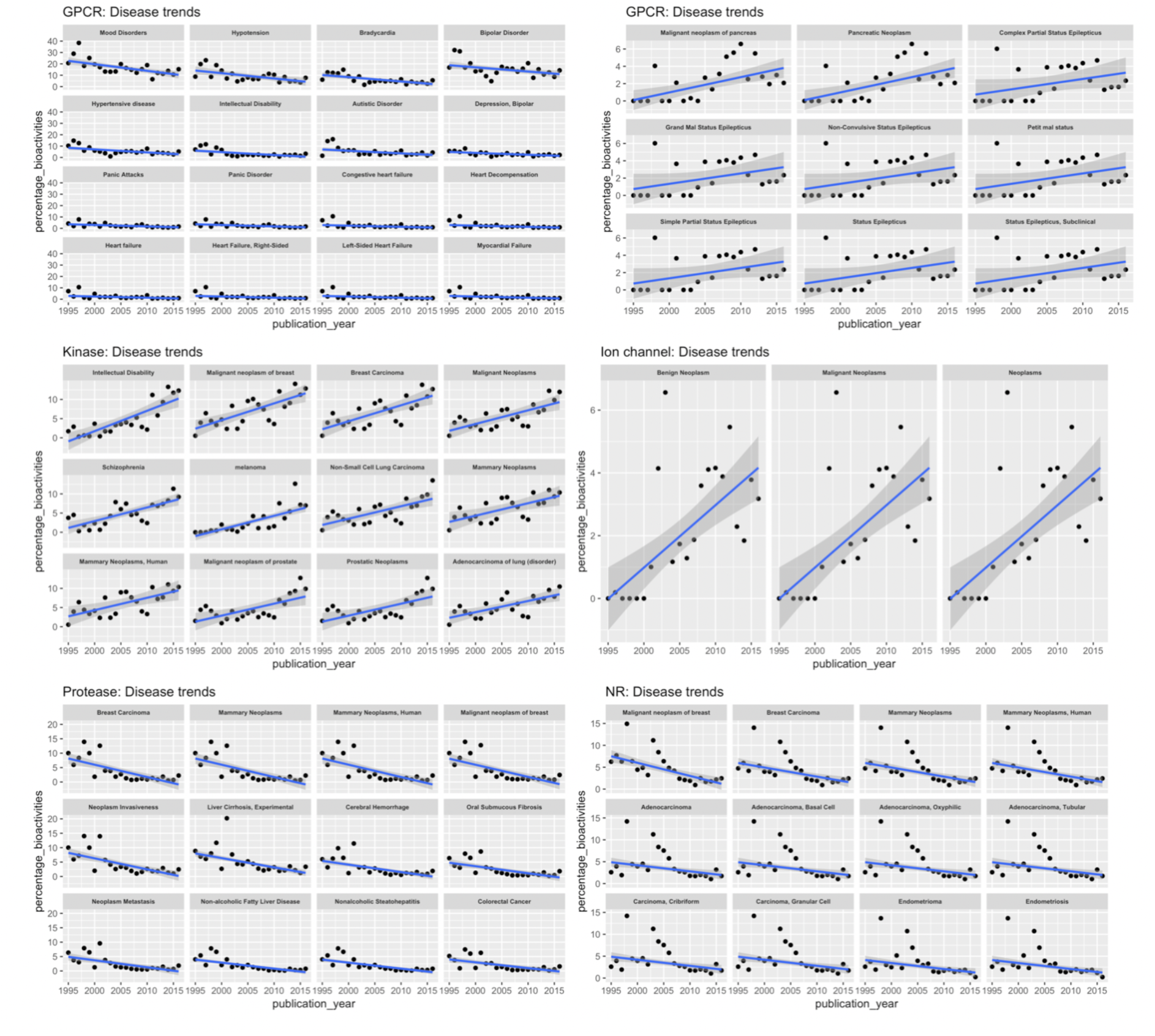
Trend line plots for disease annotations (per target family). Only the twelve steepest positive and twelve steepest negative trends are shown (if available) considering only statistically significant trends (robust regression; p <= 0.05).

**Supplementary Figure S5:**
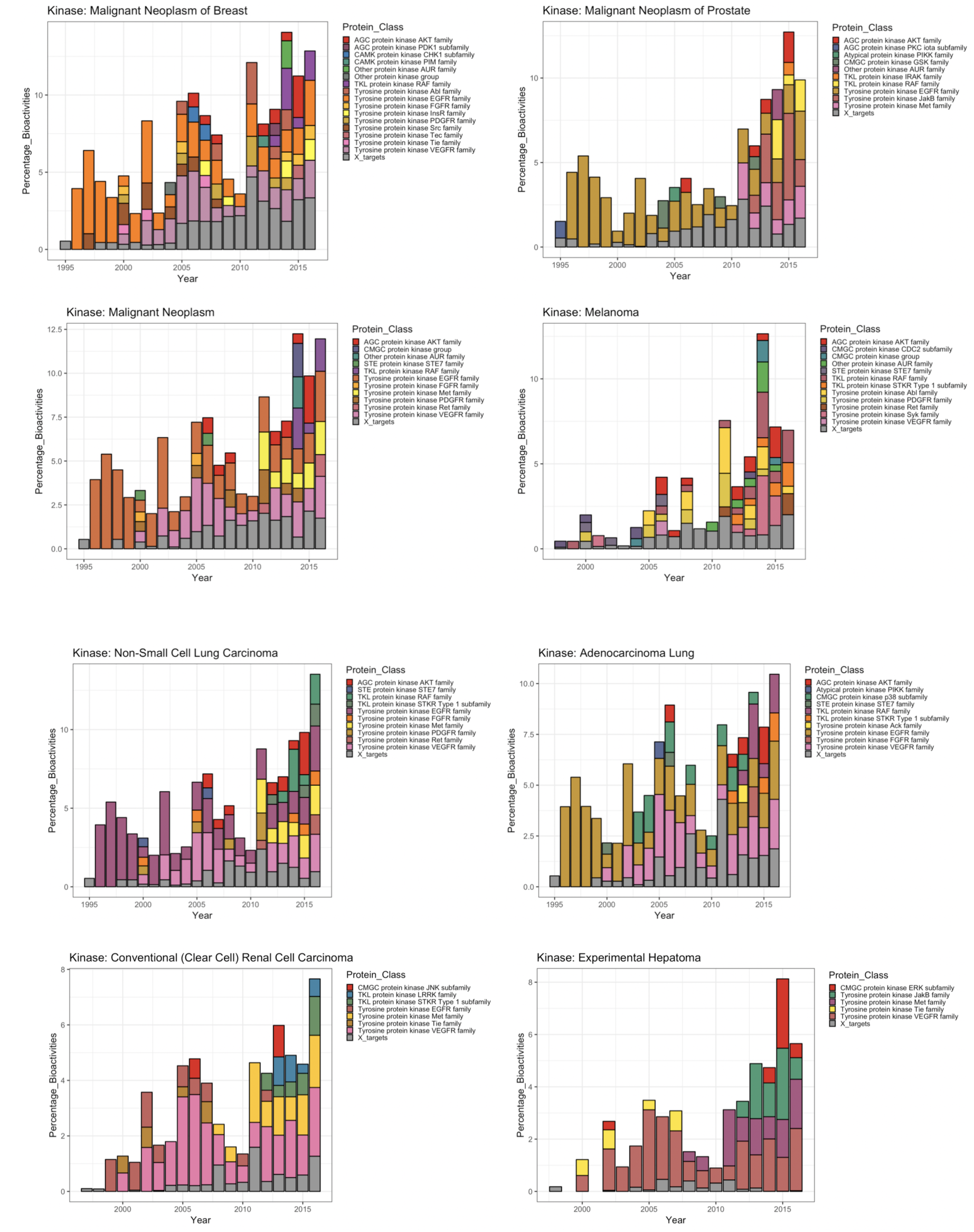
Bar charts showing the proportions of different protein classes contributing to selected steep cancer-related disease trends for kinases.

**Supplementary Figure S6:**
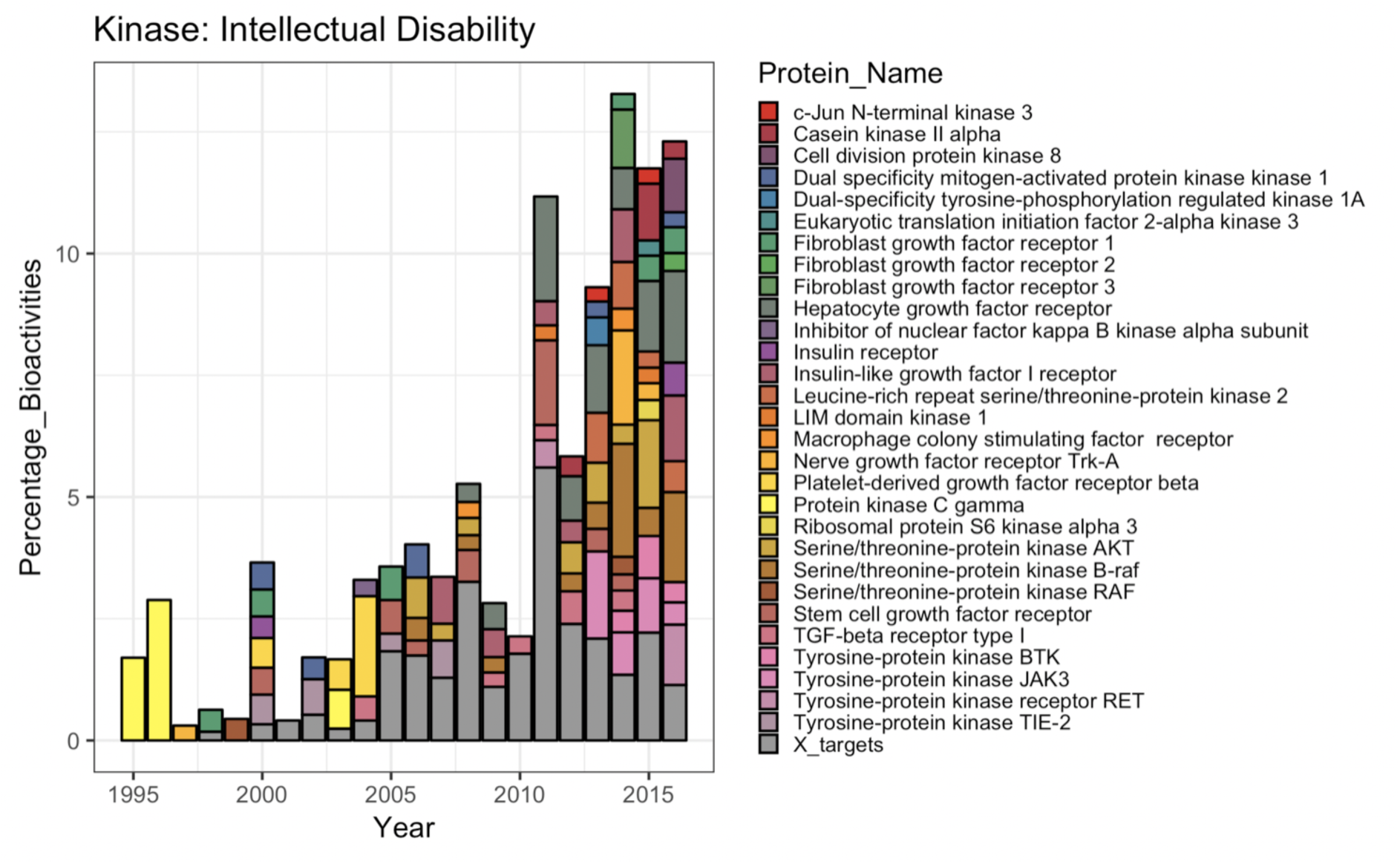
Bar chart showing the proportions of different kinases contributing to steep positive disease trend “intellectual disability”.

**Supplementary Figure S7:**
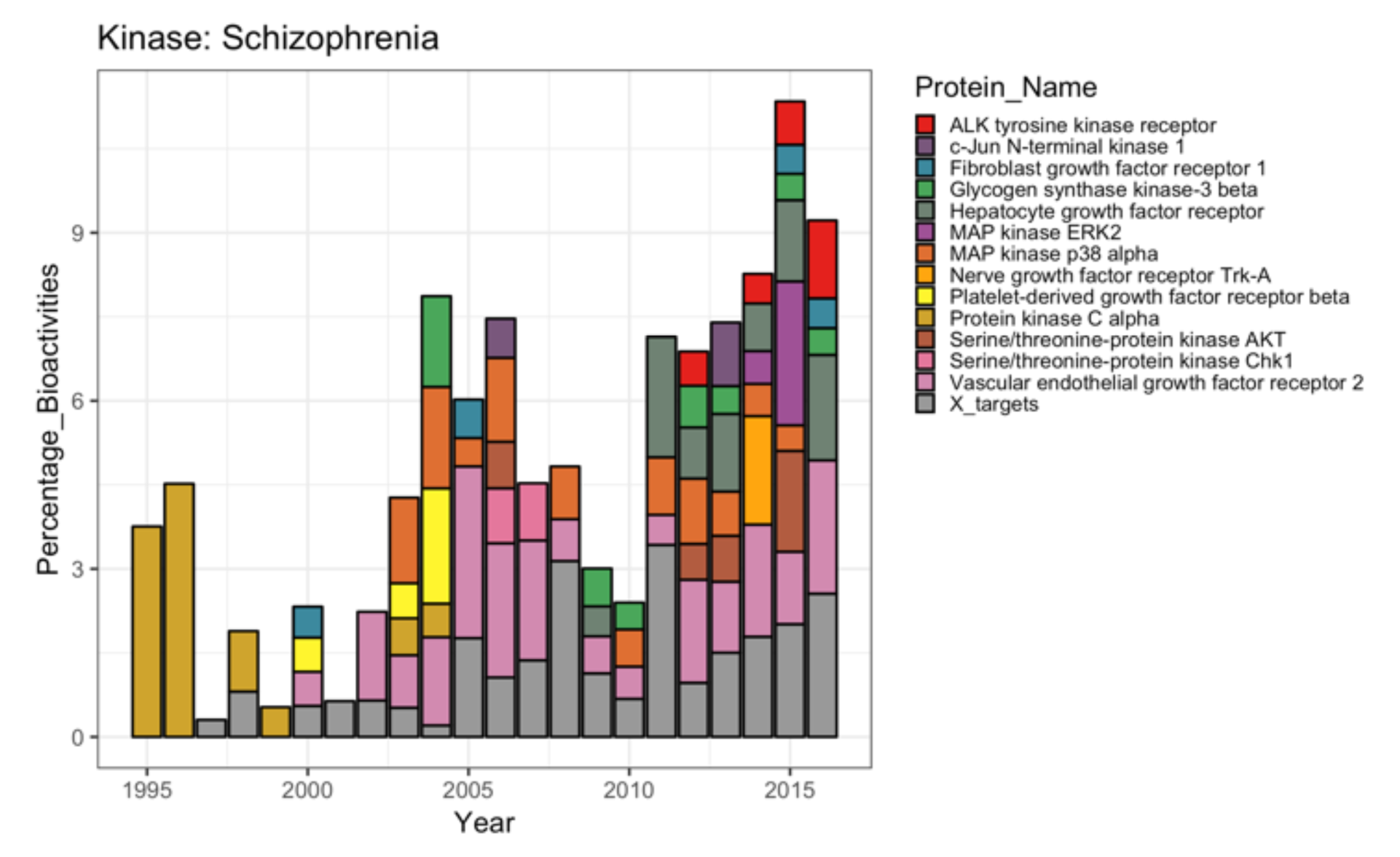
Bar chart showing the proportions of different kinases contributing to steep positive disease trend “schizophrenia”.

**Supplementary Figure S8:**
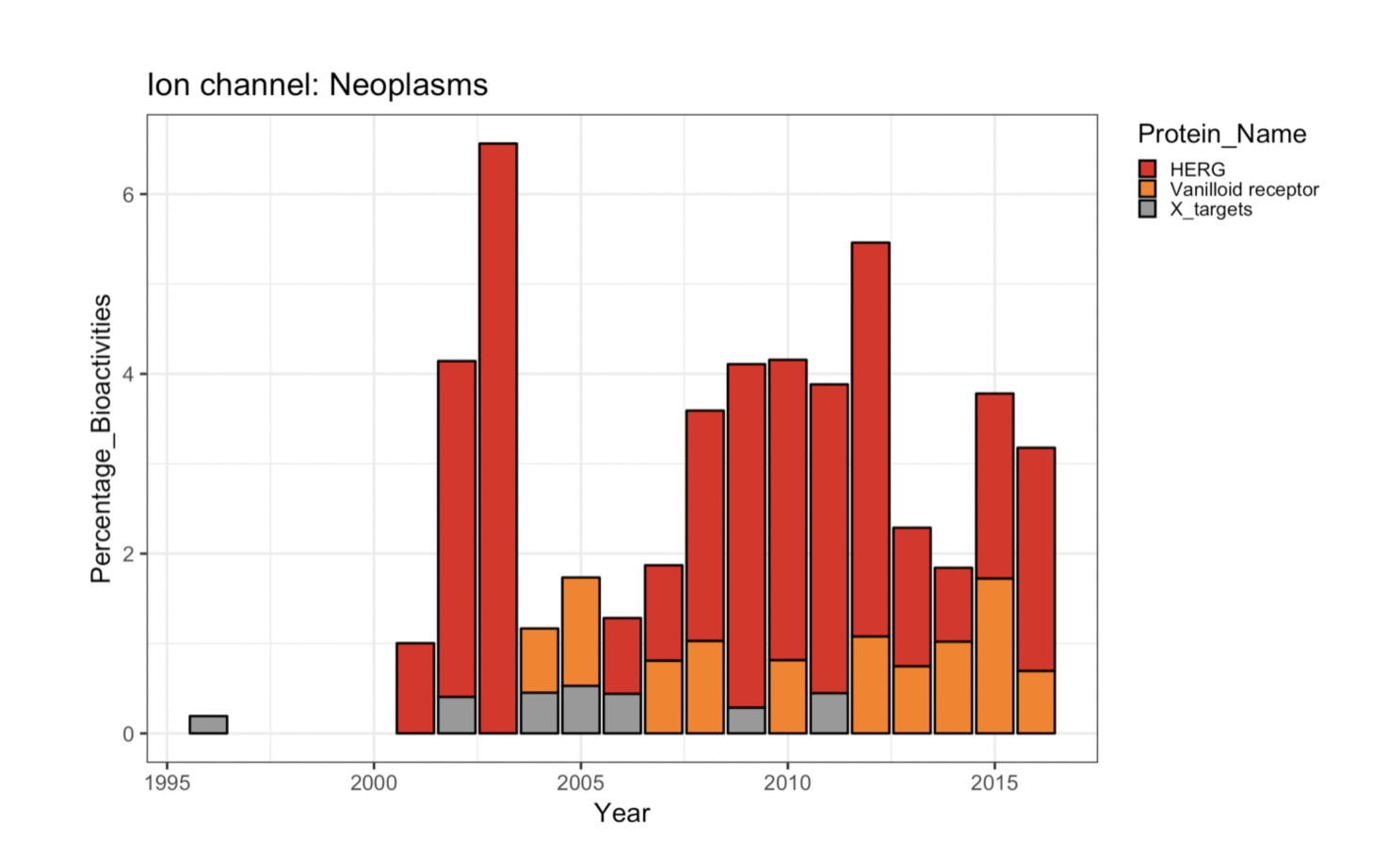
Stacked bar chart showing the proportions of different ion channels contributing to the disease trend “neoplasms”.

**Supplementary Figure S9:**
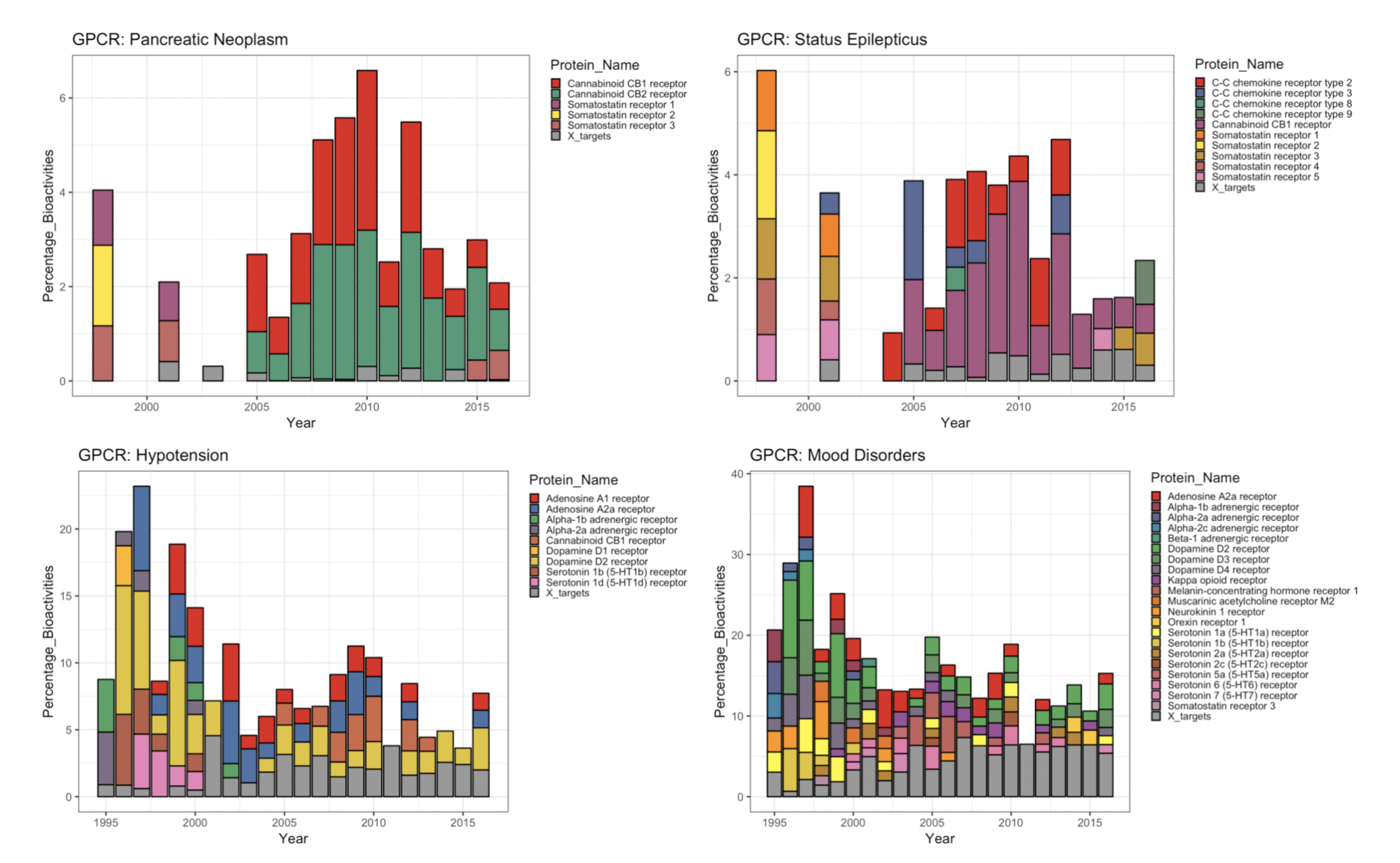
Bar charts showing the proportions of different targets contributing to steep positive and negative disease trends for GPCRs.

**Supplementary Figure S10:**
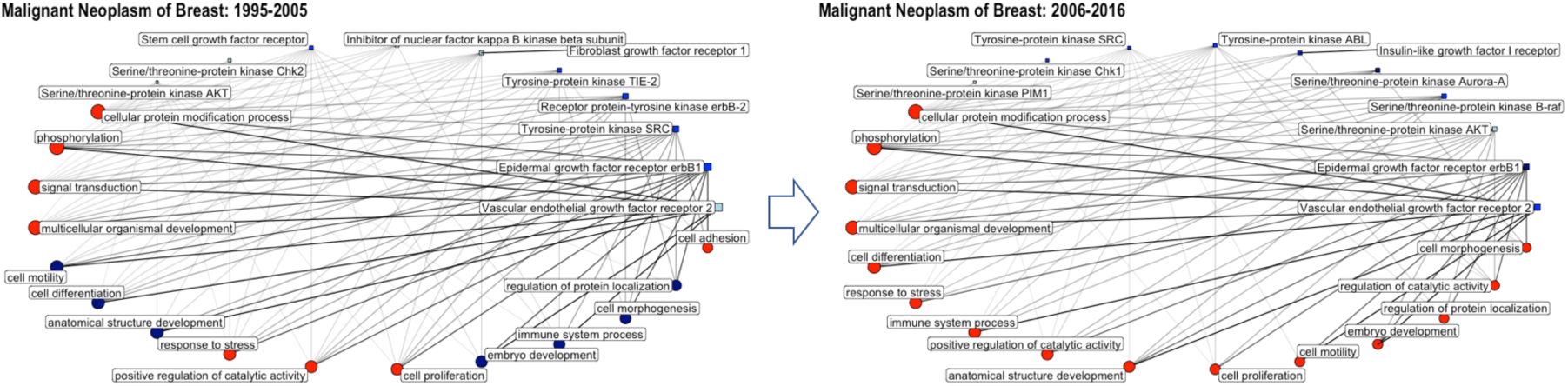
Network graphs showing the connectivity of single proteins and GO biological process annotations annotated to the disease **“malignant neoplasm of breast”** for the time periods 1995-2005 and 2006-2016. Circles correspond to GO biological process annotations; squares correspond to single protein targets; only the top 10 single protein targets and top 15 GO terms are shown (with respect to percentage of total bioactivities); the size of the circle/square as well as the thickness of the connecting lines is correlated to the percentage of bioactivities associated with this GO annotation/protein; the color code is reflecting the number of available drug-efficacy target annotations (light blue: 0; blue: 1-5; navy: 6-10; red >10).

**Supplementary Figure S11:**
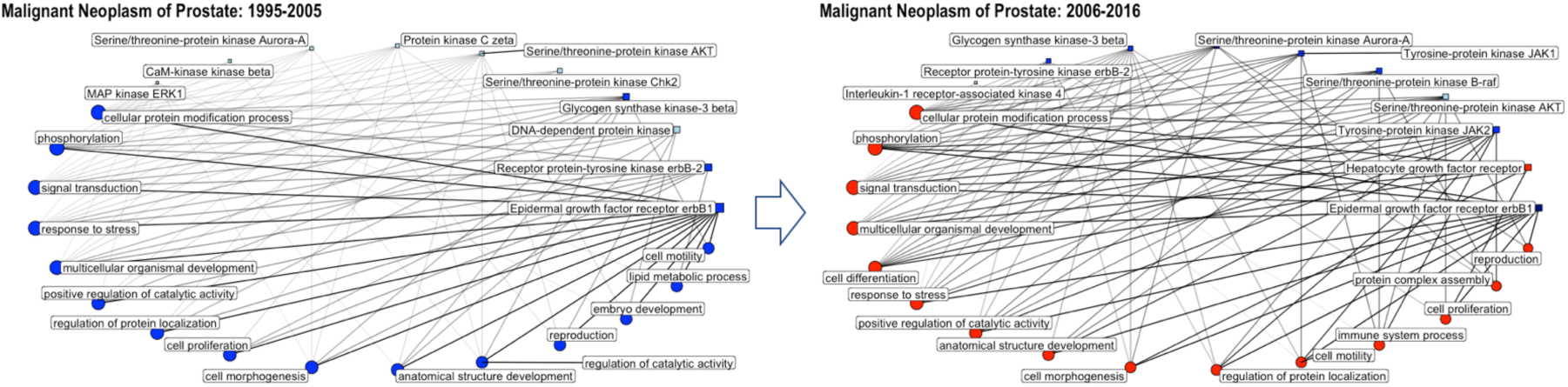
Network graphs showing the connectivity of single proteins and GO biological process annotations annotated to the disease **“malignant neoplasm of prostate”** for the time periods 1995-2005 and 2006-2016. Circles correspond to GO biological process annotations; squares correspond to single protein targets; only the top 10 single protein targets and top 15 GO terms are shown (with respect to percentage of total bioactivities); the size of the circle/square as well as the thickness of the connecting lines is correlated to the percentage of bioactivities associated with this GO annotation/protein; the color code is reflecting the number of available drug-efficacy target annotations (light blue: 0; blue: 1-5; navy: 6-10; red >10).

**Supplementary Figure S12:**
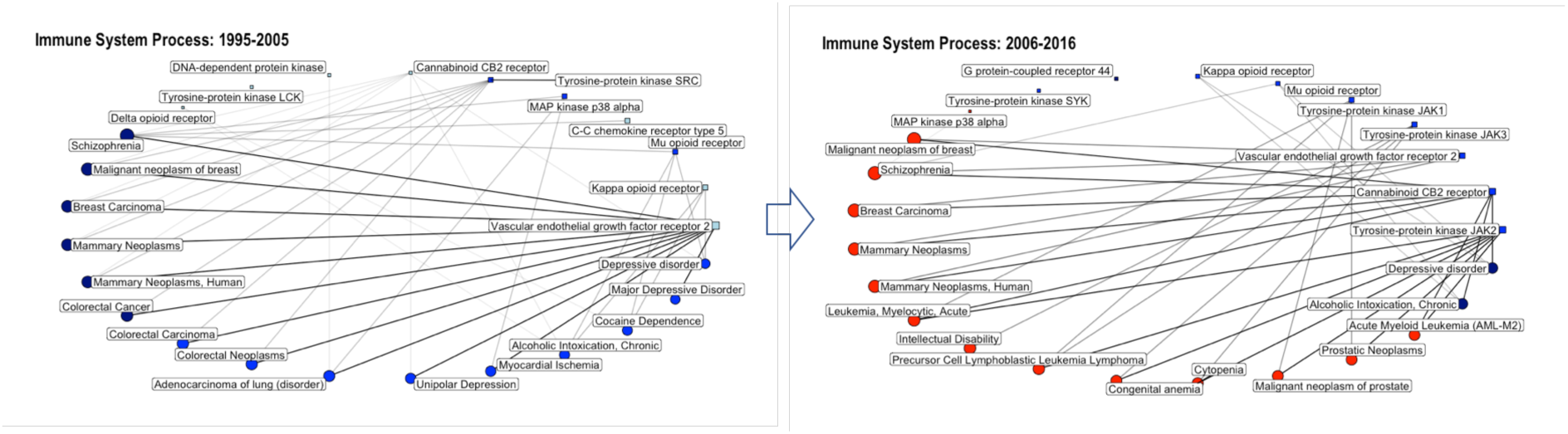
Network graphs showing the connectivity of single proteins and disease annotations annotated to the disease **“immune system process”** for the time periods 1995-2005 and 2006-2016. Circles correspond to disease annotations; squares correspond to single protein targets; only the top 10 single protein targets and top 15 disease annotations are shown (with respect to percentage of total bioactivities); the size of the circle/square as well as the thickness of the connecting lines is correlated to the percentage of bioactivities associated with this disease annotation/protein; the color code is reflecting the number of available drug-efficacy target annotations (light blue: 0; blue: 1-5; navy: 6-10; red >10).

**Supplementary Figure S13:**
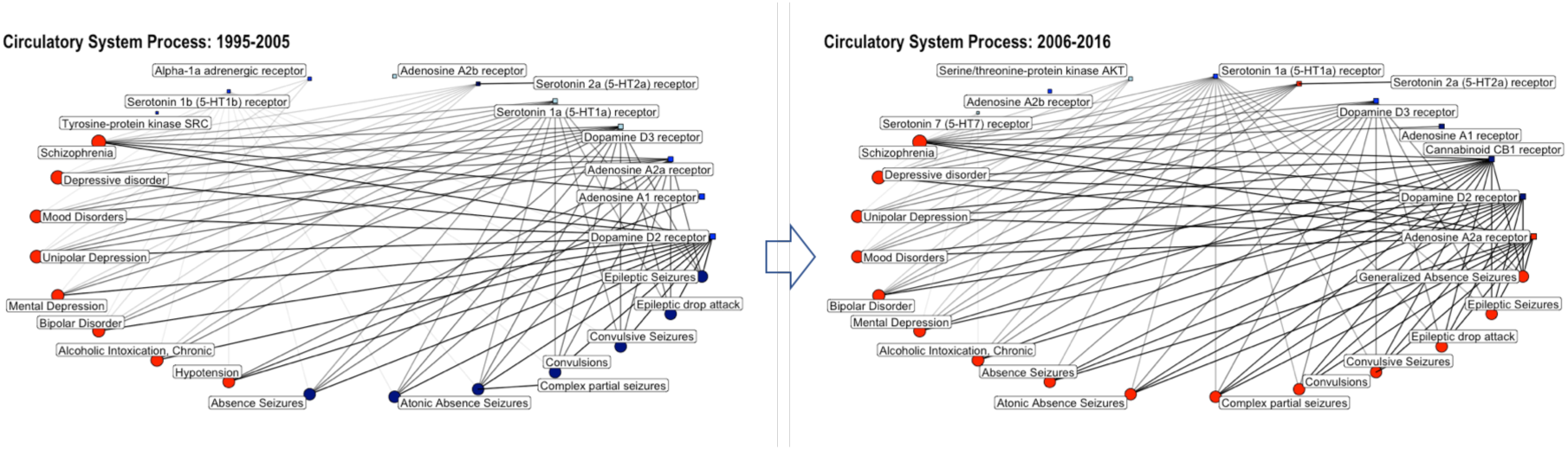
Network graphs showing the connectivity of single proteins and disease annotations annotated to the disease **“circulatory system process”** for the time periods 1995-2005 and 2006-2016. Circles correspond to disease annotations; squares correspond to single protein targets; only the top 10 single protein targets and top 15 disease annotations are shown (with respect to percentage of total bioactivities); the size of the circle/square as well as the thickness of the connecting lines is correlated to the percentage of bioactivities associated with this disease annotation/protein; the color code is reflecting the number of available drug-efficacy target annotations (light blue: 0; blue: 1-5; navy: 6-10; red >10).

**Supplementary Table S1:**
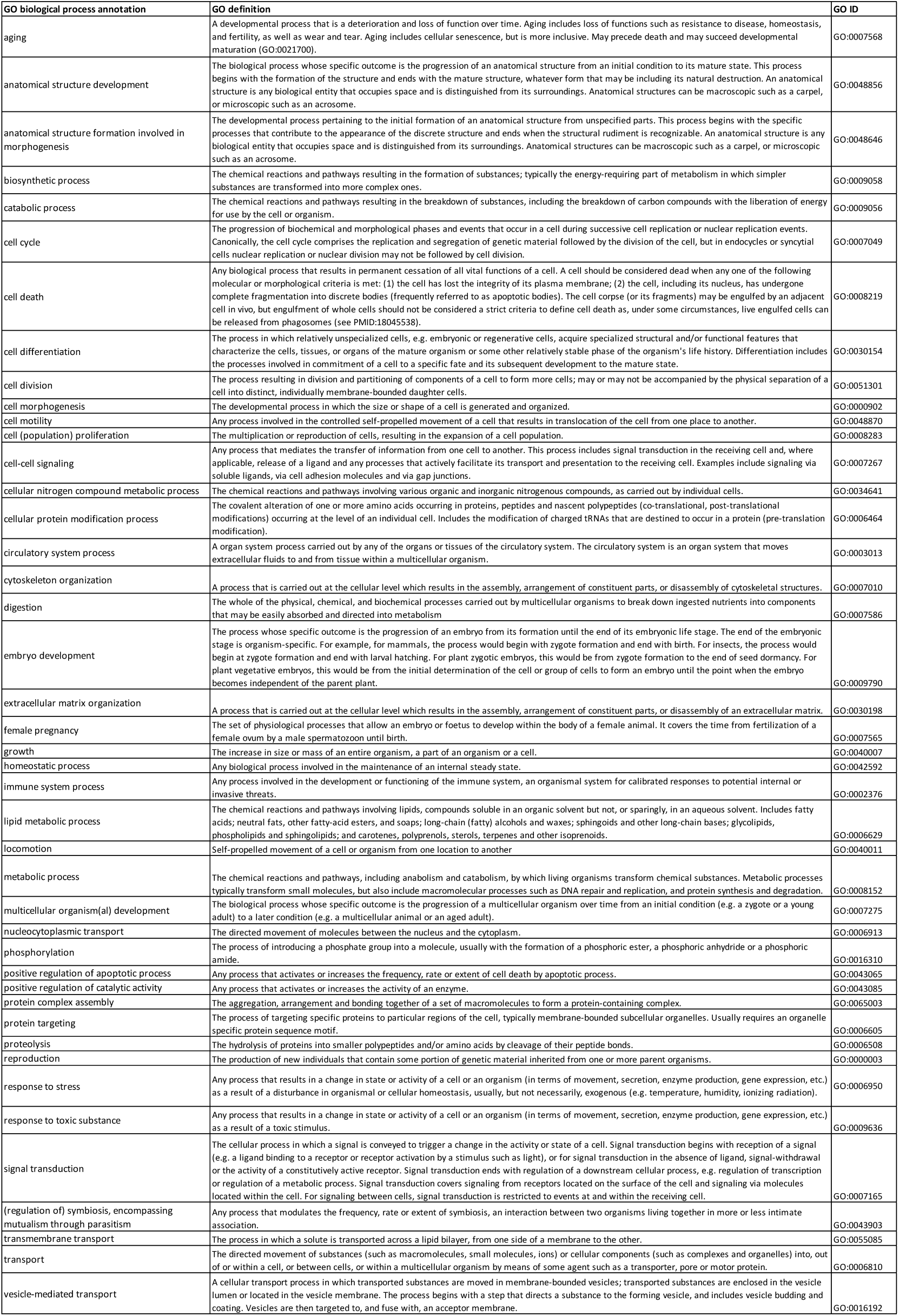
Summary of all GO biological process annotations that are annotated to the protein targets under study, including definitions for GO annotations and respective GO IDs (retrieved from https://www.ebi.ac.uk/QuickGO/ and http://www.informatics.jax.org/vocab/gene_ontology/).

**Supplementary Table 2:** Percentages and absolute numbers of bioactivities per target class for the network “Malignant neoplasm of breast” for the period 1995-2005.

| Protein Class Name | Protein Class Short Name | Percentage of Measurements | Number of Measurements |
| --- | --- | --- | --- |
| Tyrosine protein kinase EGFR family | Egfr | 1.55 | 605 |
| Tyrosine protein kinase VEGFR family | Vegfr | 1.34 | 524 |
| Tyrosine protein kinase Src family | Src | 0.58 | 226 |
| Tyrosine protein kinase FGFR family | Fgfr | 0.24 | 95 |
| Tyrosine protein kinase Tie family | Tie | 0.23 | 91 |
| Tyrosine protein kinase PDGFR family | Pdgfr | 0.20 | 80 |
| Other protein kinase group | Other | 0.19 | 74 |
| AGC protein kinase AKT family | Akt | 0.15 | 60 |
| CAMK protein kinase RAD53 family | Rad53 | 0.12 | 48 |
| Other protein kinase AUR family | Aur | 0.05 | 18 |
| Tyrosine protein kinase Abl family | Abl | 0.04 | 15 |
| CAMK protein kinase PIM family | Pim | 0.03 | 12 |
| TKL protein kinase RAF family | Raf | 0.03 | 11 |
| CAMK protein kinase CHK1 subfamily | Chk1 | 0.03 | 10 |
| Tyrosine protein kinase Eph family | Eph | 0.02 | 6 |
| CAMK protein kinase RSKb subfamily | Rskb | 0.01 | 5 |
| Tyrosine protein kinase InsR family | InsR | 0.01 | 5 |
| CAMK protein kinase AMPK subfamily | Ampk | 0.01 | 4 |
| CK1 protein kinase CK1 family | Ck1 | 0.01 | 3 |
| Tyrosine protein kinase Tec family | Tec | 0.01 | 3 |
| STE protein kinase PAKA subfamily | Paka | 0.01 | 2 |
| AGC protein kinase PDK1 subfamily | Pdk1 | 0.00 | 1 |

**Supplementary Table 3:** Percentages and absolute numbers of bioactivities per target class for the network “Malignant neoplasm of breast” for the period 2006-2016.

| Protein Class Name | Protein Class Short Name | Percentage of Measurements | Number of Measurements |
| --- | --- | --- | --- |
| Tyrosine protein kinase VEGFR family | Vegfr | 1.23 | 3926 |
| Tyrosine protein kinase EGFR family | Egfr | 1.14 | 3615 |
| AGC protein kinase AKT family | Akt | 0.70 | 2213 |
| TKL protein kinase RAF family | Raf | 0.61 | 1933 |
| Other protein kinase AUR family | Aur | 0.44 | 1407 |
| Tyrosine protein kinase InsR family | InsR | 0.40 | 1274 |
| Tyrosine protein kinase PDGFR family | Pdgfr | 0.35 | 1124 |
| Tyrosine protein kinase Abl family | Abl | 0.35 | 1123 |
| Tyrosine protein kinase Src family | Src | 0.28 | 896 |
| CAMK protein kinase CHK1 subfamily | Chk1 | 0.27 | 874 |
| Tyrosine protein kinase FGFR family | Fgfr | 0.27 | 864 |
| CAMK protein kinase PIM family | Pim | 0.25 | 809 |
| AGC protein kinase PDK1 subfamily | Pdk1 | 0.21 | 681 |
| Tyrosine protein kinase Tec family | Tec | 0.21 | 661 |
| Other protein kinase group | Other | 0.18 | 581 |
| Tyrosine protein kinase Tie family | Tie | 0.10 | 304 |
| CAMK protein kinase RSKb subfamily | Rskb | 0.07 | 223 |
| STE protein kinase PAKA subfamily | Paka | 0.06 | 203 |
| CK1 protein kinase CK1 family | Ck1 | 0.05 | 149 |
| Tyrosine protein kinase Eph family | Eph | 0.04 | 139 |
| CAMK protein kinase RAD53 family | Rad53 | 0.03 | 91 |
| TKL protein kinase STKR1 | Stkr1 | 0.02 | 65 |
| CAMK protein kinase AMPK subfamily | Ampk | 0.02 | 57 |
| Atypical protein kinase PIKK family | Pikk | 0.02 | 52 |
| STE protein kinase STE7 family | Ste7 | 0.01 | 36 |
| TKL protein kinase STKR Type 2 subfamily | Type2 | 0.01 | 29 |
| TKL protein kinase TESK subfamily | Tesk | 0.01 | 28 |
| TKL protein kinase LZK subfamily | Lzk | 0.01 | 18 |
| AGC protein kinase SGK family | Sgk | 0.01 | 17 |
| CAMK protein kinase QIK subfamily | Qik | 0.00 | 14 |
| STE protein kinase STE11 family | Ste11 | 0.00 | 12 |
| STE protein kinase ASK | Ask | 0.00 | 4 |

**Supplementary Table 4:** Percentages and absolute numbers of bioactivities per target class for the network “Malignant neoplasm of prostate” for the period 1995-2005.

| Protein Class Name | Protein Class Short Name | Percentage of Measurements | Number of Measurements |
| --- | --- | --- | --- |
| Tyrosine protein kinase EGFR family | Egfr | 1.55 | 605 |
| Atypical protein kinase PIKK family | Pikk | 0.39 | 152 |
| CMGC protein kinase GSK family | Gsk | 0.37 | 143 |
| CAMK protein kinase RAD53 family | Rad53 | 0.12 | 48 |
| AGC protein kinase AKT family | Akt | 0.10 | 41 |
| AGC protein kinase PKC iota subfamily | Iota | 0.06 | 23 |
| Other protein kinase AUR family | Aur | 0.05 | 18 |
| Other protein kinase Meta subfamily | Meta | 0.02 | 6 |
| Tyrosine protein kinase JakB family | Jakb | 0.01 | 5 |
| CMGC protein kinase ERK subfamily | Erk | 0.01 | 4 |
| TKL protein kinase STKR Type 2 subfamily | Type2 | 0.01 | 3 |
| Tyrosine protein kinase Met family | Met | 0.01 | 2 |
| STE protein kinase PAKB subfamily | Pakb | 0.00 | 1 |

**Supplementary Table 5:** Percentages and absolute numbers of bioactivities per target class for the network “Malignant neoplasm of prostate” for the period 2006-2016.

| Protein Class Name | Protein Class Short Name | Percentage of Measurements | Number of Measurements |
| --- | --- | --- | --- |
| Tyrosine protein kinase JakB family | Jakb | 1.27 | 4036 |
| Tyrosine protein kinase EGFR family | Egfr | 1.11 | 3542 |
| Tyrosine protein kinase Met family | Met | 0.79 | 2512 |
| AGC protein kinase AKT family | Akt | 0.51 | 1621 |
| TKL protein kinase RAF family | Raf | 0.51 | 1616 |
| Other protein kinase AUR family | Aur | 0.44 | 1407 |
| CMGC protein kinase GSK family | Gsk | 0.32 | 1032 |
| Atypical protein kinase PIKK family | Pikk | 0.13 | 413 |
| TKL protein kinase IRAK family | Irak | 0.09 | 280 |
| TKL protein kinase TAK1 subfamily | Tak1 | 0.04 | 126 |
| Tyrosine protein kinase DDR family | Ddr | 0.04 | 124 |
| AGC protein kinase PKC iota subfamily | Iota | 0.03 | 107 |
| Other protein kinase PKR | Pkr | 0.03 | 100 |
| CMGC protein kinase ERK subfamily | Erk | 0.03 | 94 |
| CAMK protein kinase RAD53 family | Rad53 | 0.03 | 91 |
| TKL protein kinase STKR Type 2 subfamily | Type2 | 0.02 | 73 |
| Tyrosine protein kinase FGFR family | Fgfr | 0.01 | 42 |
| TKL protein kinase STKR Type 1 subfamily | Type1 | 0.01 | 35 |
| Other protein kinase Meta subfamily | Meta | 0.01 | 34 |
| Tyrosine protein kinase Eph family | Eph | 0.01 | 34 |
| STE protein kinase PAKB subfamily | Pakb | 0.01 | 20 |
| AGC protein kinase PKA family | Pka | 0.00 | 15 |
| STE protein kinase STE11 family | Ste11 | 0.00 | 12 |

**Supplementary Table 6:** Percentages and absolute numbers of bioactivities per target class for the network “Immune system process” for the period 1995-2005.

| Protein Class Name | Protein Class Short Name | Percentage of Measurements | Number of Measurements |
| --- | --- | --- | --- |
| Opioid receptor | Opioid receptor | 1.63 | 638 |
| Tyrosine protein kinase VEGFR family | Vegfr | 1.34 | 524 |
| Tyrosine protein kinase Src family | Src | 1.13 | 444 |
| CC chemokine receptor | CC chemokine receptor | 0.87 | 342 |
| CMGC protein kinase p38 subfamily | p38 | 0.58 | 228 |
| EDG receptor | EDG receptor | 0.43 | 170 |
| Cannabinoid receptor | Cannabinoid receptor | 0.39 | 154 |
| Atypical protein kinase PIKK family | Pikk | 0.39 | 152 |
| AGC protein kinase PKC alpha subfamily | Alpha | 0.37 | 144 |
| Histamine receptor | Histamine receptor | 0.37 | 143 |
| Tyrosine protein kinase PDGFR family | Pdgfr | 0.26 | 101 |
| AGC protein kinase PKC delta subfamily | Delta | 0.25 | 99 |
| Other protein kinase group | Other | 0.25 | 97 |
| Tyrosine protein kinase Tie family | Tie | 0.23 | 91 |
| Prostanoid receptor | Prostanoid receptor | 0.22 | 87 |
| AGC protein kinase PKC eta subfamily | Eta | 0.21 | 81 |
| CXC chemokine receptor | CXC chemokine receptor | 0.16 | 63 |
| Endothelin receptor | Endothelin receptor | 0.15 | 60 |
| CAMK protein kinase MSKb subfamily | Mskb | 0.14 | 56 |
| AGC protein kinase AKT family | Akt | 0.10 | 41 |
| PAF receptor | PAF | 0.06 | 24 |
| STE protein kinase STE7 family | Ste7 | 0.06 | 23 |
| Tyrosine protein kinase Syk family | Syk | 0.05 | 20 |
| STE protein kinase STE11 family | Ste11 | 0.05 | 18 |
| TKL protein kinase STKR Type 1 subfamily | Type1 | 0.05 | 18 |
| Tyrosine protein kinase Trk family | Trk | 0.04 | 17 |
| Tyrosine protein kinase Abl family | Abl | 0.04 | 15 |
| Tyrosine protein kinase Eph family | Eph | 0.04 | 15 |
| Bradykinin receptor | Bradykinin receptor | 0.04 | 14 |
| CMGC protein kinase ERK subfamily | Erk | 0.04 | 14 |
| Leukotriene receptor | Leukotriene receptor | 0.04 | 14 |
| Anaphylatoxin receptor | Anaphylatoxin | 0.03 | 11 |
| CAMK protein kinase CAMK2 family | Camk2 | 0.03 | 11 |
| TKL protein kinase RAF family | Raf | 0.03 | 11 |
| CAMK protein kinase DAPK family | Dapk | 0.02 | 9 |
| TKL protein kinase RIPK family | Ripk | 0.02 | 9 |
| Tyrosine protein kinase JakB family | Jakb | 0.02 | 7 |
| CMGC protein kinase group | Cmgc | 0.02 | 6 |
| Tyrosine protein kinase Csk family | Csk | 0.02 | 6 |
| CAMK protein kinase RSKb subfamily | Rskb | 0.01 | 5 |
| STE protein kinase MST subfamily | Mst | 0.01 | 5 |
| Tyrosine protein kinase FGFR family | Fgfr | 0.01 | 5 |
| Tyrosine protein kinase InsR family | InsR | 0.01 | 5 |
| Tyrosine protein kinase Tec family | Tec | 0.01 | 5 |
| Tyrosine protein kinase Fak family | Fak | 0.01 | 4 |
| STE protein kinase PAKA subfamily | Paka | 0.01 | 3 |
| TKL protein kinase STKR Type 2 subfamily | Type2 | 0.01 | 3 |
| Tyrosine protein kinase Ack family | Ack | 0.01 | 3 |
| Tyrosine protein kinase Fer family | Fer | 0.01 | 3 |
| AGC protein kinase ROCK subfamily | Rock | 0.01 | 2 |
| TKL protein kinase LIMK subfamily | Limk | 0.01 | 2 |
| AGC protein kinase PDK1 subfamily | Pdk1 | 0.00 | 1 |
| Protein kinase regulatory subunit | Reg | 0.00 | 1 |

**Supplementary Table 7:** Percentages and absolute numbers of bioactivities per target class for the network “Immune system process” for the period 2006-2016.

| Protein Class Name | Protein Class Short Name | Percentage of Measurements | Number of Measurements |
| --- | --- | --- | --- |
| Opioid receptor | Opioid receptor | 2.10 | 6673 |
| Tyrosine protein kinase JakB family | Jakb | 1.92 | 6095 |
| Cannabinoid receptor | Cannabinoid receptor | 1.47 | 4663 |
| Tyrosine protein kinase VEGFR family | Vegfr | 1.23 | 3926 |
| CC chemokine receptor | CC chemokine receptor | 1.07 | 3401 |
| Prostanoid receptor | Prostanoid receptor | 0.75 | 2396 |
| Tyrosine protein kinase Syk family | Syk | 0.68 | 2147 |
| Tyrosine protein kinase Src family | Src | 0.66 | 2095 |
| Tyrosine protein kinase PDGFR family | Pdgfr | 0.63 | 2005 |
| TKL protein kinase RAF family | Raf | 0.61 | 1930 |
| EDG receptor | EDG receptor | 0.58 | 1857 |
| CMGC protein kinase p38 subfamily | p38 | 0.58 | 1838 |
| AGC protein kinase AKT family | Akt | 0.51 | 1621 |
| CXC chemokine receptor | CXC chemokine receptor | 0.51 | 1619 |
| Tyrosine protein kinase InsR family | InsR | 0.40 | 1274 |
| CMGC protein kinase ERK subfamily | Erk | 0.39 | 1233 |
| Tyrosine protein kinase Abl family | Abl | 0.35 | 1123 |
| Tyrosine protein kinase Fak family | Fak | 0.35 | 1119 |
| Tyrosine protein kinase Tec family | Tec | 0.34 | 1073 |
| Tyrosine protein kinase Trk family | Trk | 0.33 | 1065 |
| Other protein kinase group | Other | 0.26 | 838 |
| Histamine receptor | Histamine receptor | 0.26 | 828 |
| AGC protein kinase PDK1 subfamily | Pdk1 | 0.21 | 681 |
| Tyrosine protein kinase Axl family | Axl | 0.18 | 587 |
| TKL protein kinase STKR Type 1 subfamily | Type1 | 0.18 | 581 |
| Hydroxycarboxylic acid receptor | Hydroxycarboxylic acid receptor | 0.17 | 543 |
| AGC protein kinase PKC delta subfamily | Delta | 0.14 | 452 |
| STE protein kinase STE7 family | Ste7 | 0.14 | 450 |
| Tyrosine protein kinase Ack family | Ack | 0.14 | 437 |
| Tyrosine protein kinase Ret family | Ret | 0.12 | 387 |
| TKL protein kinase IRAK family | Irak | 0.11 | 363 |
| AGC protein kinase ROCK subfamily | Rock | 0.11 | 359 |
| Atypical protein kinase PIKK family | Pikk | 0.10 | 311 |
| Tyrosine protein kinase Tie family | Tie | 0.10 | 304 |
| AGC protein kinase PKC alpha subfamily | Alpha | 0.09 | 292 |
| CMGC protein kinase group | Cmgc | 0.09 | 281 |
| STE protein kinase PAKA subfamily | Paka | 0.09 | 275 |
| Tyrosine protein kinase FGFR family | Fgfr | 0.08 | 255 |
| TKL protein kinase LIMK subfamily | Limk | 0.07 | 236 |
| CAMK protein kinase RSKb subfamily | Rskb | 0.07 | 223 |
| STE protein kinase STE11 family | Ste11 | 0.07 | 207 |
| Tyrosine protein kinase Eph family | Eph | 0.06 | 175 |
| TKL protein kinase RIPK family | Ripk | 0.05 | 154 |
| Bradykinin receptor | Bradykinin receptor | 0.05 | 145 |
| CAMK protein kinase DAPK family | Dapk | 0.04 | 131 |
| Anaphylatoxin receptor | Anaphylatoxin | 0.04 | 130 |
| TKL protein kinase TAK1 subfamily | Tak1 | 0.04 | 126 |
| AGC protein kinase PKC eta subfamily | Eta | 0.03 | 110 |
| CAMK protein kinase CAMK2 family | Camk2 | 0.03 | 100 |
| Other protein kinase PKR | Pkr | 0.03 | 100 |
| CAMK protein kinase PKD family | Pkd | 0.03 | 99 |
| Leukotriene receptor | Leukotriene receptor | 0.03 | 88 |
| STE protein kinase MST subfamily | Mst | 0.03 | 82 |
| CMGC protein kinase HIPK subfamily | Hipk | 0.03 | 80 |
| Tyrosine protein kinase Csk family | Csk | 0.02 | 76 |
| TKL protein kinase STKR Type 2 subfamily | Type2 | 0.02 | 73 |
| CAMK protein kinase MELK subfamily | Melk | 0.02 | 67 |
| Tyrosine protein kinase Met family | Met | 0.02 | 66 |
| AGC protein kinase PKA family | Pka | 0.02 | 64 |
| Protease-activated receptor | Protease-activated receptor | 0.02 | 58 |
| Other protein kinase GCN2 subfamily | Gcn2 | 0.02 | 54 |
| Purine receptor | Purine receptor | 0.02 | 52 |
| CX3C chemokine receptor | CX3C chemokine receptor | 0.02 | 51 |
| Other protein kinase IKK family | Ikk | 0.01 | 46 |
| AGC protein kinase PKN family | Pkn | 0.01 | 43 |
| CAMK protein kinase MSKb subfamily | Mskb | 0.01 | 36 |
| Other protein kinase HRI | Hri | 0.01 | 34 |
| Tyrosine protein kinase Fer family | Fer | 0.01 | 33 |
| STE protein kinase unique family | Ste-Unique | 0.01 | 31 |
| Tyrosine protein kinase Srm | Srm | 0.01 | 30 |
| Family A G protein-coupled receptor | 7TM1 | 0.01 | 22 |
| CAMK protein kinase LKB subfamily | Lkb | 0.01 | 20 |
| CMGC protein kinase SRPK family | Srpk | 0.01 | 19 |
| CAMK protein kinase CAMK1 family | Camk1 | 0.01 | 18 |
| Endothelin receptor | Endothelin receptor | 0.00 | 12 |
| CAMK protein kinase MAPKAPK subfamily | Mapkapk | 0.00 | 7 |
| PAF receptor | PAF | 0.00 | 4 |
| Steroid-like ligand receptor | Steroid-like ligand receptor | 0.00 | 4 |

**Supplementary Table 8:** Percentages and absolute numbers of bioactivities per target class for the network “Circulatory system process” for the period 1995-2005.

| Protein Class Name | Protein Class Short Name | Percentage of Measurements | Number of Measurements |
| --- | --- | --- | --- |
| Serotonin receptor | Serotonin receptor | 3.51 | 1373 |
| Adenosine receptor | Adenosine receptor | 3.18 | 1247 |
| Adrenergic receptor | Adrenergic receptor | 2.95 | 1155 |
| Dopamine receptor | Dopamine receptor | 2.65 | 1039 |
| Neurokinin receptor | Neurokinin receptor | 0.85 | 334 |
| Vasopressin and oxytocin receptor | Vasopressin and oxytocin receptor | 0.80 | 312 |
| Tyrosine protein kinase Src family | Src | 0.64 | 249 |
| Cannabinoid receptor | Cannabinoid receptor | 0.54 | 213 |
| Melatonin receptor | Melatonin receptor | 0.42 | 164 |
| Endothelin receptor | Endothelin receptor | 0.40 | 155 |
| Opioid receptor | Opioid receptor | 0.39 | 151 |
| Histamine receptor | Histamine receptor | 0.37 | 143 |
| Acetylcholine receptor | Acetylcholine receptor | 0.34 | 134 |
| Melanocortin receptor | Melanocortin receptor | 0.33 | 128 |
| Prostanoid receptor | Prostanoid receptor | 0.32 | 124 |
| Protease-activated receptor | Protease-activated receptor | 0.27 | 104 |
| Tyrosine protein kinase Tie family | Tie | 0.23 | 91 |
| Bradykinin receptor | Bradykinin receptor | 0.19 | 76 |
| Neuropeptide Y receptor | Neuropeptide Y receptor | 0.11 | 44 |
| AGC protein kinase AKT family | Akt | 0.10 | 41 |
| Neuropeptide receptor | Neuropeptide receptor | 0.08 | 30 |
| Purine receptor | Purine receptor | 0.07 | 27 |
| PAF receptor | PAF | 0.06 | 24 |
| Angiotensin receptor | Angiotensin receptor | 0.05 | 19 |
| Leukotriene receptor | Leukotriene receptor | 0.03 | 12 |
| CAMK protein kinase RSKb subfamily | Rskb | 0.02 | 7 |
| CAMK protein kinase MLCK family | MLck | 0.01 | 4 |
| CMGC protein kinase GSK family | Gsk | 0.01 | 4 |
| Atypical protein kinase BCR family | Bcr | 0.01 | 3 |
| AGC protein kinase PKC delta subfamily | Delta | 0.01 | 2 |
| CAMK protein kinase CAMK2 family | Camk2 | 0.01 | 2 |
| AGC protein kinase SGK family | Sgk | 0.00 | 1 |
| CXC chemokine receptor | CXC chemokine receptor | 0.00 | 1 |

**Supplementary Table 9:** Percentages and absolute numbers of bioactivities per target class for the network “Circulatory system process” for the period 2006-2016.

| Protein Class Name | Protein Class Short Name | Percentage of Measurements | Number of Measurements |
| --- | --- | --- | --- |
| Adenosine receptor | Adenosine receptor | 2.57 | 8169 |
| Serotonin receptor | Serotonin receptor | 2.19 | 6951 |
| Dopamine receptor | Dopamine receptor | 2.16 | 6857 |
| Adrenergic receptor | Adrenergic receptor | 1.34 | 4267 |
| Cannabinoid receptor | Cannabinoid receptor | 1.09 | 3473 |
| Acetylcholine receptor | Acetylcholine receptor | 0.56 | 1768 |
| AGC protein kinase AKT family | Akt | 0.51 | 1621 |
| Neurokinin receptor | Neurokinin receptor | 0.45 | 1441 |
| Vasopressin and oxytocin receptor | Vasopressin and oxytocin receptor | 0.38 | 1194 |
| Tyrosine protein kinase Src family | Src | 0.32 | 1028 |
| Opioid receptor | Opioid receptor | 0.31 | 970 |
| Histamine receptor | Histamine receptor | 0.26 | 828 |
| Bradykinin receptor | Bradykinin receptor | 0.24 | 766 |
| Prostanoid receptor | Prostanoid receptor | 0.22 | 713 |
| Melatonin receptor | Melatonin receptor | 0.15 | 483 |
| Angiotensin receptor | Angiotensin receptor | 0.12 | 391 |
| Protease-activated receptor | Protease-activated receptor | 0.12 | 377 |
| Melanocortin receptor | Melanocortin receptor | 0.11 | 351 |
| Purine receptor | Purine receptor | 0.11 | 341 |
| Neuropeptide receptor | Neuropeptide receptor | 0.11 | 338 |
| Tyrosine protein kinase Tie family | Tie | 0.10 | 304 |
| AGC protein kinase PKC delta subfamily | Delta | 0.09 | 276 |
| Neuropeptide Y receptor | Neuropeptide Y receptor | 0.06 | 206 |
| CXC chemokine receptor | CXC chemokine receptor | 0.06 | 204 |
| Leukotriene receptor | Leukotriene receptor | 0.05 | 173 |
| AGC protein kinase SGK family | Sgk | 0.03 | 97 |
| CMGC protein kinase GSK family | Gsk | 0.03 | 95 |
| Neurotensin receptor | Neurotensin receptor | 0.03 | 93 |
| CAMK protein kinase RSKb subfamily | Rskb | 0.03 | 84 |
| Endothelin receptor | Endothelin receptor | 0.02 | 78 |
| CAMK protein kinase MLCK family | Mlck | 0.02 | 54 |
| STE protein kinase STE7 family | Ste7 | 0.02 | 49 |
| AGC protein kinase BARK subfamily | Bark | 0.01 | 41 |
| TKL protein kinase STKR Type 1 subfamily | Type1 | 0.01 | 37 |
| CAMK protein kinase CAMK2 family | Camk2 | 0.01 | 30 |
| TKL protein kinase STKR Type 2 subfamily | Type2 | 0.01 | 29 |
| AGC protein kinase PKG family | Pkg | 0.00 | 14 |
| STE protein kinase FRAY subfamily | Fray | 0.00 | 12 |
| PAF receptor | PAF | 0.00 | 4 |
| Steroid-like ligand receptor | Steroid-like ligand receptor | 0.00 | 4 |

